# Ultrahigh yields of giant vesicles obtained through mesophase evolution and breakup

**DOI:** 10.1101/2024.06.03.597257

**Authors:** Alexis Cooper, Anand Bala Subramaniam

## Abstract

Self-assembly odry amphiphilic lipid films on surfaces upon hydration is a crucial step in the formation of cell-like giant unilamellar vesicles (GUVs). GUVs are useful as model biophysical systems, as chassis for synthetic biology, and in biomedical applications. Here via combined quantitative measurements of the molar yield and distributions of sizes and high-resolution imaging of the evolution of lipid films on surfaces, we report the discovery of a previously unknown pathway for the assembly of GUVs which can lead to ultrahigh yields of > 50 %. This yield is about 60 % higher than any GUV yield reported to date. The “shear-induced fragmentation” pathway occurs in membranes containing 3 mol % of the poly(ethylene glycol) modified lipid PEG2000-DSPE (1,2-distearoyl-*sn*-glycero-3-phosphoethanolamine-N-[methoxy(polyethylene glycol)-2000]), when a lipid-dense foam-like mesophase forms upon hydration. The membranes in the mesophase fragment and close to form GUVs upon application of fluid shear. Experiments with varying mol % of PEG2000-DSPE and with lipids with partial molecular similarity to PEG2000-DSPE show that ultrahigh yields are only achievable under conditions where the lipid-dense mesophase forms. The increased yield of GUVs compared to mixtures without PEG2000-DSPE was general to other flat supporting surfaces such as stainless steel sheets and to various lipid mixtures. Since FDA-approved liposomal and lipid nanoparticle formulations use PEG2000-DSPE, these results provide a useful route to obtaining ultrahigh yields of GUVs that are suitable for biomedical applications.

## Introduction

Giant unilamellar vesicles (GUVs), single-walled phospholipid vesicles with diameters ≥ 1 µm, are useful for biophysical experiments since they mimic key properties of cellular membranes in an experimentally tractable minimal model.^1–3^ Additionally, GUVs show promise for use in biomedical applications^4–8^ and for bottom-up synthetic biology.^9–11^

GUVs are routinely obtained by hydrating dry thin lipid films with low ionic strength aqueous solutions,^12–15^ yet the pathways of assembly of GUVs remains to be fully understood. Recent experimental advances have allowed the interrogation of the pathways of assembly of GUVs upon hydration of thin lipid films. By correlating the evolution of the film on the surface with quantitative measurements of the yields and the distribution of sizes of populations of GUVs, we studied the effects of physicochemical parameters such as the geometry and chemical composition of the substrate used to support the lipid film^13^, the lipid composition^12^, the identity of assisting polymeric compounds,^12^ the duration^16^ and the temperature^12^ of incubation, and the ionic strength of the hydrating solution^12^. Depending on the conditions, GUV yields ranged from low, 0.3 – 4.9 %, moderate, 5 – 19.9 %, and high, 20 – 40 %. Surfaces composed of enmeshed nanocylinders, such as the nanocellulose paper used in the PAPYRUS method, Paper-Abetted amPhiphile hYdRation in aqUeous Solutions, resulted in a higher yield of GUVs compared to the flat surfaces used in the gentle hydration and electroformation methods.^13,16^

Mechanistically, comparisons of the free energy of the lipids configured as a spherical bud versus a bilayer remaining on the substrate showed that the free energy change to form a bud can be negative for bilayers on nanocylinders while the free energy change to form a bud is always positive for bilayers on flat surfaces. Once formed, merging of the buds with each other to reduce the total number of buds reduces the elastic energy of the system. A hypothetical pathway involving both budding and merging provides an energetically favorable path for bilayers on nanoscale cylinders to form large GUV-sized buds. In contrast, the path on flat surfaces requires the input of energy^13^. A similar free energy argument shows that the dissolution of partially soluble polymers exerts an osmotic pressure that can balance the increased adhesion between bilayers due to the screening of electrostatic charges in salty solutions. The osmotic pressure contribution lowers the free energy of budding, thus resulting in high yields of GUVs in salty solutions^12^.

In this work, we investigate the effects of adding 0.1-10 mol % of poly(ethylene glycol) modified (PEGylated) lipids on the assembly pathway of GUVs composed of the zwitterionic lipid DOPC (1,2-dioleoyl-*sn*-glycero-3-phosphocholine) through the PAPYRUS and gentle hydration methods. PEG2000-DSPE (1,2-distearoyl-*sn*-glycero-3-phosphoethanolamine-N-[methoxy(polyethylene glycol)-2000]) has a long history of use in clinical lipid formulations because it improves the circulation lifetime by reducing opsonization of liposomes and lipid nanoparticles by serum proteins^17–32^. The lipid consists of an uncharged hydrophilic polymer of 45 repeating ethylene glycol monomer units with an average molecular weight of 2000 g/mol that is covalently attached to the headgroup of the phospholipid DSPE. The utility of PEG2000-DPSE for biomedical applications has led to many experimental and theoretical investigations of its effects on the physicochemical properties of membranes such as intermembrane forces^33–36^, phase behavior^37–40^, mechanical property^39,41–45^, permeability^44,46–48^, and protein adsorption. Model systems that have been investigated include oriented multilayers^32,35,36^, supported lipid bilayers^32,35,49^, small unilamellar liposomes^40,46,47^, GUVs^44^, multilamellar vesicles^33,34,36,38^, and Langmuir monolayers^50^. Many studies involve lipid membranes in the gel-phase^32–35,38,40,44,47,50,51^. Comparatively less is known about the effects of PEGylated lipids on the pathway of assembly of GUVs^52,53^ or in fluid phase membranes^36,46^, despite the wide use of GUVs with fluid phase membranes in biophysical experiments^1–3,9,54^.

We find, unexpectedly given previous results ^13,16^, that we can obtain yields of > 50 % of GUVs when the lipid mixture contained 3 mol % PEG2000-DSPE using the gentle hydration method on flat glass surfaces in low salt solutions. This yield is ∼ 175 % higher than the yield of GUVs obtained using pure DOPC through the gentle hydration method. The yield is the highest that has been measured thus far, leading us to add a classification of “ultrahigh” for yields > 45 %. Interestingly, obtaining this ultrahigh yield was highly specific to the gentle hydration method and only when the mixture had 3 mol % PEG2000-DSPE. We find that there was no difference in GUV yields between the mixture containing 3 mol % PEG2000-DSPE and pure DOPC obtained using the PAPYRUS method. Furthermore, performing the gentle hydration method using mixtures with other mol percents of PEG2000-DSPE resulted in yields that ranged from 16 – 30 %.

High resolution three-dimensional confocal microscopy images of the hydrated lipid films on the surface show that the DOPC + 3 mol % PEG2000-DSPE lipid mixture rapidly forms a lipid-dense foam-like mesophase. Although a qualitatively similar mesophase forms with a DOPC + 5 mol % PEG2000-DSPE lipid mixture, the lipid content of the mesophase was low and micelles were present in the solution. There was no mesophase for the other mol percents of PEG2000-DSPE or on the surface of nanocellulose paper. Instead, the lipid assembles into surface-attached buds for mixtures containing ≤ 1 mol % of PEG2000-DSPE and sparse free-floating clusters of GUVs and micelles for mixtures containing 10 mol % PEG2000-DSPE. Importantly, we find that different configurations of the lipid on the surface can lead to similar yields of GUVs upon harvesting.

We use lipids with partial molecular similarity to PEG2000-DSPE to understand the physical interactions that promote the formation of the foam-like mesophase. We find that incorporation of lipids that do not have both a charged headgroup and a covalently attached PEG2000 chain did not result in the formation of the foam-like mesophase nor did it result in GUV yields different from pure DOPC. Additionally, we find that screening of the charge on PEG2000-DSPE by using solutions with dissolved salts prevents the formation of the foam-like mesophase and results in a dramatic drop in GUV yields. We also show that an increase in GUV yield upon addition of 3 mol % PEG2000-DSPE appears general to other lipid mixtures and other flat surfaces. Guided by these results, we propose a mechanism of assembly of GUVs using lipid mixtures containing 3 to 5 mol % PEG2000-DSPE. We propose that during harvesting with a pipette, the lamellar membranes in the foam-like mesophase fragments due to fluid shear and self-close to form GUVs. This shear-induced breakup of the mesophase is an alternate, previously unrecognized, pathway for forming GUV with fluid phase membranes containing PEG2000-DSPE that can lead to ultrahigh yields under specific conditions.

## Materials and Methods

### Chemicals

We purchased sucrose (BioXtra grade, purity ≥ 99.5%), glucose (BioXtra grade, purity ≥ 99.5%), sodium chloride (BioXtra grade, purity ≥ 99.5%) and casein from bovine milk (BioReagent grade) from Sigma-Aldrich (St. Louis, MO). We purchased chloroform (ACS grade, purity ≥ 99.8%, with 0.75% ethanol as preservative) Thermo Fisher Scientific (Waltham, MA). We obtained 18.2 MΩ ultrapure water from a Milli-Q® IQ 7000 Ultrapure Lab Water System (Burlington, MA). We purchased 1,2-dioleoyl-*sn*-glycero-3-phosphocholine (DOPC), 1,2-dipalmitoyl-sn-glycero-3-phosphocholine (DPPC), cholesterol (ovine wool), 1,2-dioleoyl-sn-glycero-3-phosphoethanolamine-N-(lissamine rhodamine B sulfonyl) (Rhod-PE), 23-(dipyrrometheneboron difluoride)-24-norcholesterol (TopFluor-Chol), 1,2-distearoyl-*sn*-glycero-3-phosphoethanolamine-N-[methoxy(polyethylene glycol)-2000] (PEG2000-DSPE), distearoyl-rac-glycerol-PEG2K (PEG2000-DSG), 1,2-distearoyl-*sn*-glycero-3-phospho-(1’-rac-glycerol) (sodium salt) (DSPG), 1,2-distearoyl-sn-glycero-3-phosphoethanolamine (DSPE) from Avanti Polar Lipids, Inc. (Alabaster, AL).

### Materials

We purchased premium plain glass microscope slides (75 mm × 25 mm, catalog number: 12-544-1) from Thermo Fisher Scientific (Waltham, MA). We purchased a hole punch cutter (EK Tools) and acid free artist grade tracing paper from (Jack Richeson and Co. Inc) on Amazon. We purchased 0.001 inch (0.025 mm) thick stainless steel sheets from McMaster-Carr.

### Cleaning of substrates

We clean all substrates as previously described^13^. We clean glass slides by sequentially sonicating for 10 minutes in acetone, 200 proof ethanol, and ultrapure water. We then dry the slides under a stream of ultrapure nitrogen to remove any visible water. We allow the slides to dry further in an oven set to 65 °C for a minimum of 2 hours.

We cut sheets of 0.001 inch (0.025 mm) thick stainless steel into rectangles of 75 mm × 25 mm using a glass slide as a template. We clean the stainless steel rectangles following the same procedure for cleaning glass slides.

We clean the tracing paper as previously described^13^ by soaking the tracing paper in 100 mL of chloroform in a 500 mL glass beaker for 30 minutes with occasional manual agitation. We discard the chloroform and repeat the same process with fresh chloroform. We remove the tracing paper and leave the paper in the fume hood for 30 minutes. The substrates are then soaked in 500 mL of ultrapure water for 30 minutes. We discard the ultrapure water and repeat the process with fresh ultrapure water. The tracing paper is then placed on a clean sheet of aluminum foil and dried in a 65°C oven for 2 hours.

### Preparation of lipid stocks

Lipid mixtures were prepared as previously described with minor adaptations^12,13,16^. We created working solutions of DOPC with 0, 0.1, 1, 3, or 10 mol % PEG2000-DSPE and 0.5 mol % TopFluor-Chol. We also created working solutions of DOPC with 3 mol % each of DSPE, PEG2000-DSG, or DSPG and 0.5 mol % TopFluor-Chol. The DOPC mol % was adjusted to accommodate the minority lipids. For phase separating GUVs, we created working solutions of DOPC/DPPC/Cholesterol/PEG2000-DSPE/Rhodamine-DOPE/TopFluor-Chol at 34.5/34.5/27.5/3/0.25/0.25 mol %, and DOPC/DPPC/Cholesterol/Rhodamine-DOPE/TopFluor-Chol at 36/36/27.5/0.25/0.25 mol %. All working solutions had a concentration of 1 mg/mL. The lipid solutions were stored in clean glass vials with Teflon lined caps, purged with argon to avoid lipid oxidation, and stored in a -20 °C freezer. Lipid solutions were used within 7 days of preparation.

### Deposition of lipids

To ensure that the nominal surface concentration remained constant, we deposited 10 µL of the lipid working solution onto circles with a diameter of 9.5 mm. For application on glass slides, we punched three 3/8-inch diameter holes on the adhesive side of a 3 × 3 in Post-it^®^ using a circular punch (EK Tools Circle Punch, 3/8 in.). The Post-it^®^ was then affixed to the underside of the glass slide to create a removable and reusable template for spreading the lipid within a 9.6 mm diameter circular area. For assembly on tracing paper, we punched out a 9.6 mm diameter circular piece of tracing paper using a clean circular punch. For assembly on stainless steel sheets, we cut a template from paper using a circular punch and cut the stainless steel sheets to size using a pair of scissors. Lipids were deposited using a glass Hamilton syringe. Then, the lipid-coated substrates were placed in a standard laboratory vacuum desiccator for 1 hour to remove traces of chloroform.

### Procedure for GUV assembly

For the gentle hydration method, circular poly(dimethyl)siloxane (PDMS) gaskets (inner diameter × height = 12 × 1 mm) were affixed to the glass slides around the dried lipid films to create a hydration chamber. We then added 150 µL of a solution of 100 mM sucrose and allowed the lipid films to hydrate for 1 hour at room temperature, 22 °C. We do not place a coverslip on the chamber. To reduce evaporation, we enclosed the samples and a water-saturated Kimwipe in a 150 mm diameter Petri dish. The films were allowed to hydrate for 1 h.

We assemble phase-separated GUVs on a hotplate set to 45 °C to be above the transition temperature of DPPC (41 °C). A similar experimental setup to the procedure for assembly at room temperature resulted in the solution evaporating completely within 1 hour. We thus modified the hydration procedure by stacking three circular PDMS gaskets to create a tall hydration chamber (inner diameter × height = 12 × 3 mm) which allowed us to place a coverslip on the chamber without contacting the liquid. We found that when the coverslip contacted the liquid, removal of the coverslip results in fluid shear that affected reproducibility. We placed the sample in a closed 150 mm diameter Petri dish and set the dish on the hotplate for 1 minute to preheat. An average of ∼ 17 µL of liquid evaporates from the chamber. To account for evaporation, we add 167 µL of a solution of 90 mM sucrose. To solution was preheated to 45 °C before adding it into the hydration chamber. We then gently placed a coverslip on top of the tall PDMS chamber, placed a water-saturated Kimwipe into the dish, and replaced the lid on the Petri dish. The films were allowed to hydrate. After 1 h we removed the coverslips carefully to avoid the condensed water droplets from falling into the sample. We measured the osmolarity of a 20 µL aliquot using a freezing point depression multi-sample osmometer (Model 2020, Advanced Instruments, USA). Using this procedure, we obtained an average final sample volume of 150 ± 3 µL with an osmolarity of 100 ± 2 mM.

For the PAPYRUS method and gentle hydration method using the stainless steel sheets, the 9.5 mm diameter circles of the lipid-coated tracing paper or stainless steel sheets were placed in individual wells of a 48-well plate. 150 µL of a solution of 100 mM sucrose was then added to the wells and incubated for 1 hour at room temperature.

The GUVs were harvested from all the surfaces by pipetting 100 µL of the solution up and down 6 times with a micropipette. On the 7^th^ time, we fully aspirate the solution containing the GUVs and store it in an Eppendorf tube. Aliquots were taken immediately for imaging. Each condition was performed N=3 independent times.

### Imaging of the harvested vesicles

Imaging was conducted as previously described^13^. Briefly, we imaged the GUVs in custom-made square poly(dimethylsiloxane) (PDMS) chambers with dimensions of 5 mm × 5 mm × 1 mm (length × width × height) affixed to a glass slide. We passivated the chambers with 1 mg/mL casein dissolved in PBS. We sedimented a 2 µL aliquot of the sucrose-filled GUVs in 58 µL of an isomolar solution of glucose in the imaging chamber. The GUVs are allowed to sediment for 3 hours, followed by imaging using a confocal laser scanning microscope (LSM 880, Axio Imager.Z2m, Zeiss, Germany), using a 488 nm argon laser and a 10× Plan-Apochromat objective with a numerical aperture of 0.45. For phase separated vesicles, we capture dual channel images using a 488 nm argon laser and a 561 nm diode pumped solid state laser. The intensity and gain of the two channels were adjusted so that the mean intensity of both channels was similar. We imaged using an automated tile-scan with an autofocus routine (49 images [5951.35 µm × 5951.35 µm, (3212 pixels x 3212 pixels)]) to capture the entire area of the chamber.

### Image processing and analysis

We processed and analyzed the images as previously described.^13^ We use a custom MATLAB (Mathworks Inc., Natick, MA) routine to segment fluorescent objects from the background and the *regionprops* function to obtain the diameters and pixel intensities of the objects. We calculate the coefficient of variance (CV) of the intensities to select GUVs from the detected fluorescent objects. All images were inspected after automated segmentation and errors in segmentation and selection were corrected manually.

### Statistical Tests

All statistical tests were performed in MATLAB. We conducted a one-way balanced analysis of variance (ANOVA) when comparing more than two mean yields. These were for the mean yields obtained with 0-10 mol % of PEG2000-DSPE and for samples with 3 mol % DSPE, PEG2000-DSG or DSPG. For tests that showed a significant difference between the means, we conducted a posthoc Tukey’s honestly significant difference (HSD) to determine the statistical significance between pairs of means. We conduct Student’s *t-*tests to determine the statistical significance of the difference in mean yields when only two sample means were compared.

### High resolution Z-Stacks

We took confocal Z-Stacks using a Plan-Apochromat 20× DIC M27 75mm water dipping objective with a numerical aperture of 1.0. We took 189 slices of the hydrated lipid films at intervals of 0.9 µm starting from 4.5 µm into the surface of the substrate to 165.6 µm above the surface of the substrate. The images were 151.82 × 151.82 µm (1272 × 1272 pixels). Imaging locations were chosen to best represent the surface. Image processing was performed in FIJI^55^. We used the reslice function to create *x-z* projections of the confocal Z-Stacks. We equalize the histogram in FIJI with a pixel saturation of 0.3% to show dim features along with bright features. We obtain Z-projections by summing the slices of the first 18 µm, starting 3 slices below the surface. The histograms of the Z-projections were not modified.

## Results and Discussion

### The addition of 3 mol % PEG2000-DSPE to the lipid mixture affects the yield and distribution of sizes differently depending on the method of assembly

We assembled GUVs with membrane compositions of 99.5:0.5 mol % DOPC:TopFluor®-Chol and 97.5:3:0.5 mol % DOPC:PEG2000-DSPE:TopFluor®-Chol using the PAPYRUS and gentle hydration methods^13^. For brevity, we refer to these lipid compositions as “pure DOPC” and “DOPC + 3 mol % PEG2000-DSPE” respectively. TopFluor®-Chol is a fluorescent sterol used to visualize the lipid membranes through confocal fluorescence microscopy. Figure 1a shows the molar yields of GUVs obtained using the PAPYRUS method and Figure 1b shows the molar yield obtained using the gentle hydration method. Each bar is an average of N=3 independent repeats and the error bars are one standard deviation from the mean. Following previous convention^12,13,16^, we divide the molar yield into GUVs with diameters, *d*, between 1 µm ≤ *d* < 10 µm, 10 µm ≤ *d* < 50 µm, and *d* ≥ 50 µm. The population classes were chosen because GUVs between 1 µm ≤ *d* < 10 µm are of the sizes of blood cells, intracellular organelles, and bacteria, and GUVs between 10 µm ≤ *d* < 50 µm are of the size of mammalian cells. We show representative images of the harvested GUVs and the histogram of diameters in Figure S1 and Figure S2.

**Figure 1.**
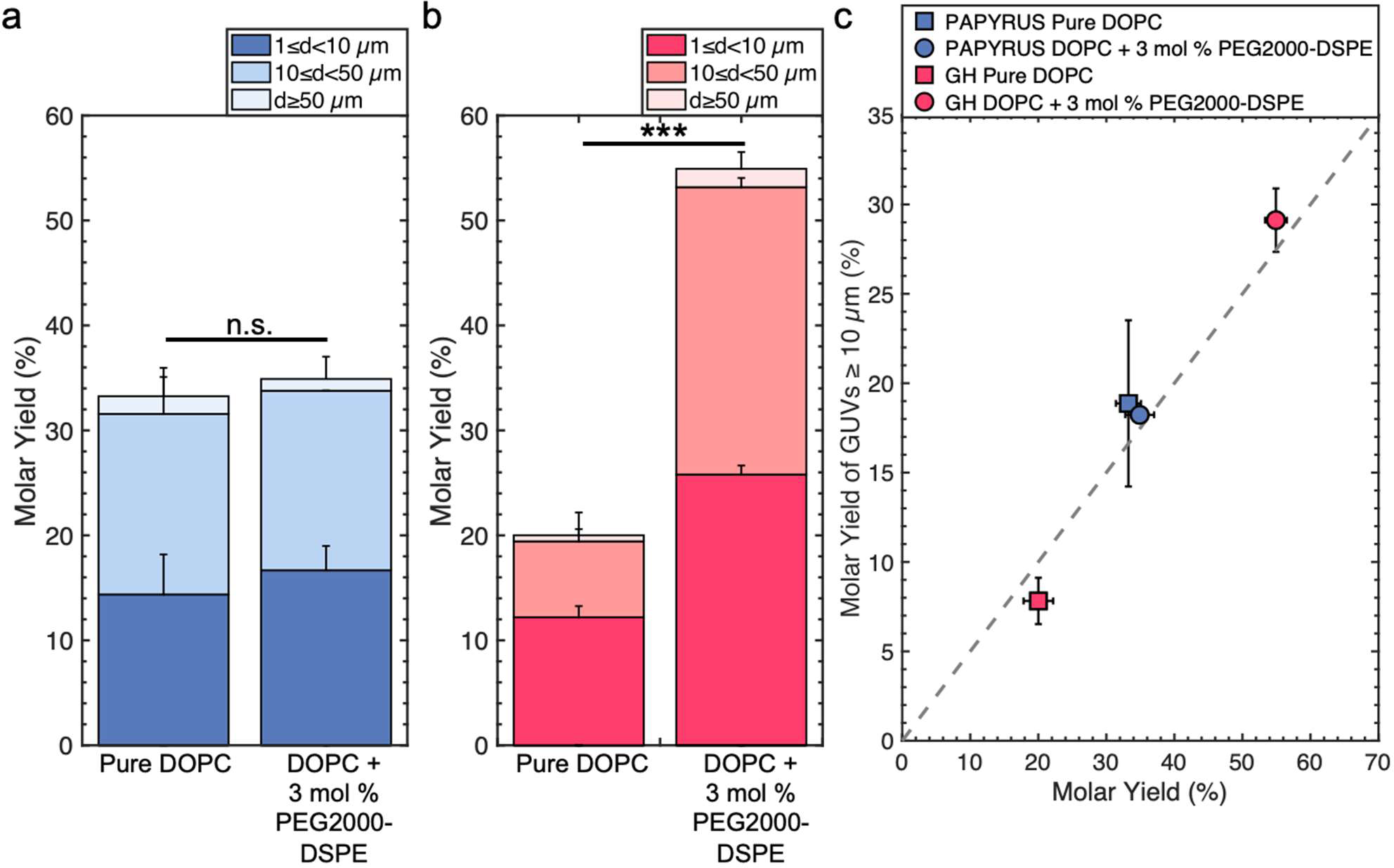
Molar yields of GUVs from mixtures of pure DOPC and DOPC + 3 mol % PEG2000-DSPE. a) Stacked bar plots showing the molar yield of GUVs obtained from the PAPYRUS method. b) Stacked bar plots showing the molar yield of GUVs obtained from the gentle hydration method. The stacks show the percentage of the molar yield that is comprised of the size classifications as listed in the legend. c) Scatter plot showing the molar yield of GUVs with diameters ≥ 10 µm versus the total molar yield. The grey dashed line indicates where half the yield is in GUVs with diameters ≥ 10 µm. Each data point is the average of N=3 independent repeats. The error bars are one standard deviation from the mean. Statistical significance was determined using a Student’s *t*-test. *: p < 0.05, **: p < 0.01, ***: p < 0.001, n.s.: not significant.

The mean yield of GUVs obtained using the PAPYRUS method is 33 ± 2 % and 35 ± 2 % for pure DOPC and DOPC + 3 mol % PEG2000-DSPE respectively (Fig. 1a). The 2 % difference in the mean yields between these two lipid compositions was not statistically significant (*p*=0.451). The mean yield of GUVs obtained using the gentle hydration method is 20 ± 2 % and 55 ± 2 % for pure DOPC and DOPC + 3 mol % PEG2000-DSPE respectively (Fig. 1b). The 35 % difference in yield between the two lipid compositions was highly statistically significant (*p* = 5.22 × 10^-5^). Furthermore, the ∼ 20 % difference in yield between DOPC + 3 mol % PEG2000-DSPE obtained using the gentle hydration method compared to the yields obtained using the PAPYRUS method for both compositions of lipid was highly statistically significant (both *p* < 0.001). Indeed, the 55 % yield is the highest yield that has been measured to date for any thin film hydration method^12,13,16^. To qualitatively distinguish the yield that we report in this work with the “high” yields reported previously, which ranged from 20-40 %,^12,13,16^ we term molar yields above 45 % as “ultrahigh” yields.

We plot the yield of GUVs with *d* ≥ 10 µm versus the total molar yield of GUVs to show the effects of the addition of 3 mol % PEG2000-DSPE on the distribution of sizes (Figure 1c). The gray dashed line represents the boundary where half of the molar yield is from GUVs with *d* ≥ 10 µm. Consistent with previous results, GUVs with *d* ≥ 10 µm make up to ∼57 % and 39 % of the total yield for the lipid composition consisting of pure DOPC obtained using the PAPYRUS and gentle hydration methods^13,16^. In contrast, GUVs with *d* ≥ 10 µm make up to 52 % and 53 % of the total yield for the lipid composition consisting of DOPC + 3 mol % PEG2000-DSPE obtained using the PAPYRUS and gentle hydration methods.

These results show that the addition of 3 mol % PEG2000-DSPE to the lipid mixtures affects the yield and distribution of sizes differently depending on the method of assembly. For the PAPYRUS method, the addition of 3 mol % of PEG2000-DSPE to the lipid mixture does not affect the yield but decreases the fraction of GUVs with *d* ≥ 10 µm. For the gentle hydration method, the addition of 3 mol % PEG2000-DSPE to the lipid mixture increases both the yield and fraction of GUVs with *d* ≥ 10 µm.

We evaluate these results within the framework of the budding and merging (BNM) model for GUV assembly^13,16^. Equation 1 and 2 gives the free energy for the formation of spherical buds of radius *R*_B_ from cylindrical bilayers, Δ*E*_RB,e_, and disk-shaped flat bilayers, Δ*E*_RB,d_.

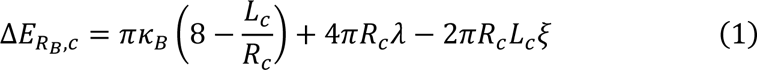

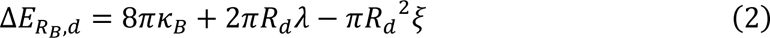

In these equations, the bending modulus, *K_B_*, the edge energy, λ, and the effective adhesive contact potential, ζ, are the physical parameters of the lipid bilayer. The radius of the cylinder, *R_e_*, the length of the cylinder, *L_C_*, and the radius of the disk, *R_d_*, are the geometrical parameters of the bilayer. The first term on the right-hand side measures the change in the bending energy to change the curvature of the bilayer, the second term measures the change in the edge energy if breaks in the bilayer must form to allow budding at a constant area, and the third term measures the change in adhesion energy to separate the bilayers. We take that the bilayers are in a stack of multiple bilayers. Thus, the effective adhesive contact potential, ζ, is that of bilayers interacting with each other.

The free energy change for forming a spherical bud from a cylindrical bilayer can be negative while the free energy change for forming a spherical bud from a flat bilayer is always positive^13,16^. This can be readily shown by using characteristic dimensions of a cylindrical bilayer on a nanocellulose fiber, *R_e_* = 20 nm, *L*_e_ = 2000 nm and a flat disk *R*_D_ = 242 nm, and with *K*_B_ = 8.5 × 10^-20^ J, λ = 1 × 10^-11^J m^-1^, and ζ = −1 × 10^-5^ J m^-2^ for DOPC.^56^ The energy to form a bud of radius *R*_B_= 141 nm from the cylindrical bilayer is Δ*E*_Rb,e_ ≈ −4754 k_B_T and from the flat bilayer is Δ*E*_Rb,d_ ≈ 5439 k_B_T. Here the energy is expressed relative to the thermal energy scale 1 k_B_T = 4.11 × 10^-21^ J. Because of the large and positive free energy change for the formation of buds from flat bilayers relative to the thermal energy scale, the formation of most GUV-sized buds occurs within 1 minute of hydration due to energy released upon hydration.^16^ For the PAPYRUS method, in addition to bud formation upon hydration, additional nanoscale buds form and merge up to 30 minutes post-hydration.^16^ Merging of the nanoscale buds with each other and with GUV-sized buds results in a higher molar yield and a greater fraction of GUVs with large diameters compared to the gentle hydration method. The budding and merging model is consistent with the assembly of GUVs using pure DOPC.

We consider changes to the physical constants of the membrane which could explain the effect of incorporation of PEG2000-DSPE on the yield of GUVs. To explain the observed results, there should be no change in the magnitude of the free energy for bud formation for budding from cylindrical bilayers whereas the magnitude of the free energy change for bud formation for budding from flat disk-shaped bilayers should be greatly reduced. Keeping the same geometrical parameters, a reduction of the magnitude of *K*_B_ by five times to 1.7 × 10^-2^ J results in Δ*E*_R,e_ = 27.5 k_B_T and Δ*E*_Rb,d_ = 5023 k_B_T. Taking the limit of no adhesion between bilayers, ζ = 0, Δ*E*_Rb,e_ = −5367 k_B_T and Δ*E*_Rb,d_ = 4831 k_B_T. Taking the limit of no edge energy, λ = 0, Δ*E*_Rb,e_ = −5366 k_B_T and Δ*E*_Rb,d_ = 1128 k_B_T. We surmise that, i) a change in the bending modulus can have a large effect on the free energy change of budding from cylindrical bilayers but has a small effect on the free energy change of budding from flat bilayers, ii) a change in the adhesive contact potential results in similar changes in the magnitudes of the free energy change of budding from cylindrical bilayers and flat bilayers, and iii) a change in the edge energy has a small effect on the free energy change of budding from cylindrical bilayers but can have a large effect on the free energy change of budding from flat bilayers.

Since changes in the physical parameters of the membrane do not provide a conclusive explanation for our results, we hypothesized that an additional pathway of assembly other than budding and merging may be operational in membranes containing 3 mol % of PEG2000-DSPE.

### Films of DOPC+3 mol % PEG2000-DSPE form a foam-like mesophase on glass but not on nanocellulose paper

To test our hypothesis, we probed directly by imaging the lipid films 1 hour after hydration using high resolution confocal microscopy. We show representative orthogonal *x - z* slices in the upper panels and *x - y* sum projections in the lower panels of Figure 2a-d. We equalized the histogram of the *x-z* slices to show both bright and dim regions. We show original images without histogram equalization in Figure S3. The other images in Figure 2 retain their original intensity values.

**Figure 2.**
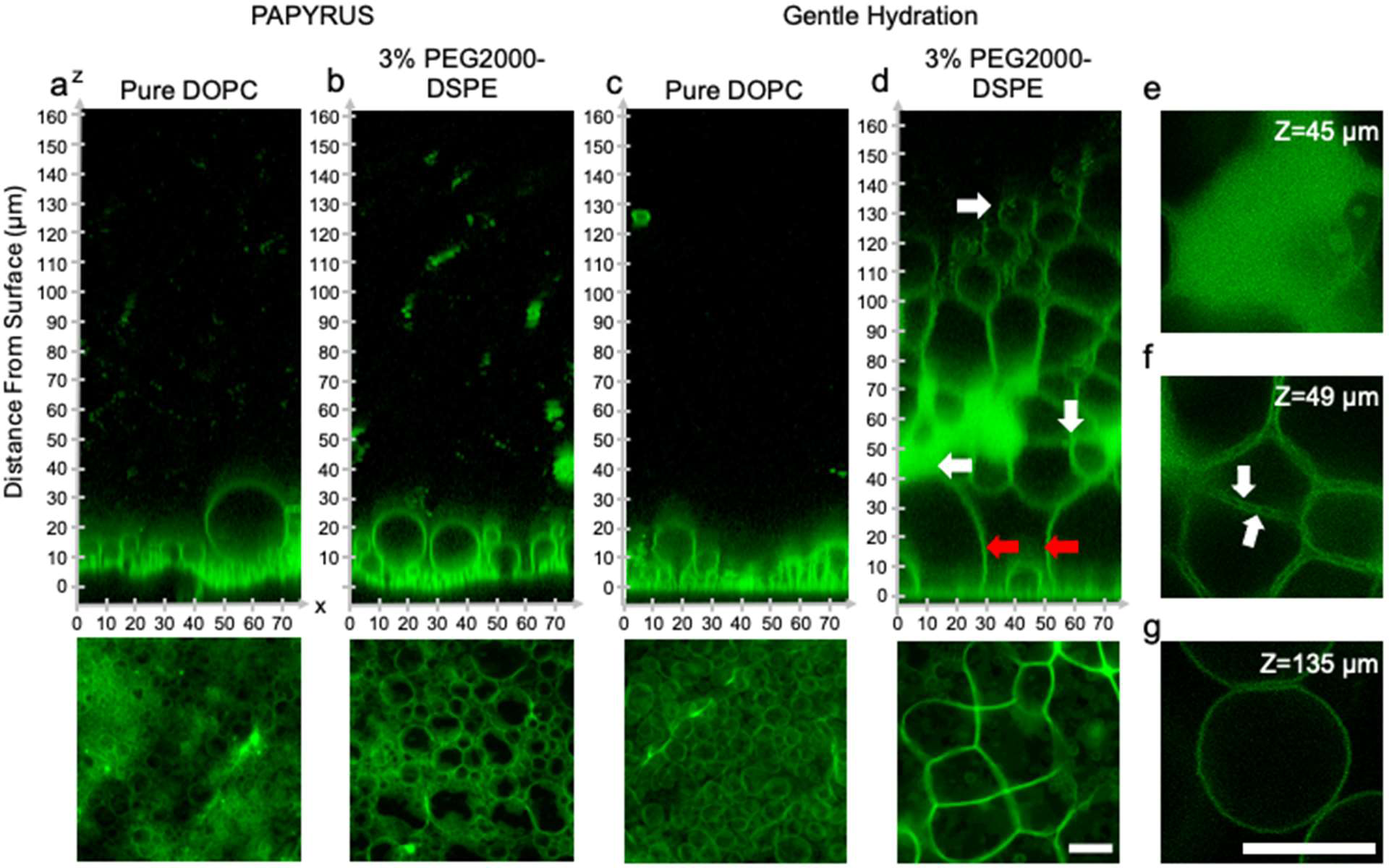
Configuration of the hydrated lipid films containing pure DOPC and DOPC + 3 mol % PEG2000-DSPE. a-d) Upper panels show orthogonal *x-z* planes of confocal Z-Stacks of the lipid films on the surface. Bottom panels are *x-y* planes consisting of summed z-projections of the first 18 µm from the surface of the substrate. a) Pure DOPC on nanocellulose paper. b) DOPC + 3 mol % PEG2000-DSPE on nanocellulose paper. c) Pure DOPC on glass. d) DOPC + 3 mol % PEG2000-DSPE on glass. The thick red arrows show the lipid septa in the mesophase. e-g) *x-y* slices from the regions indicated by the thick white arrows in d). Scale bars are 15 µm.

Similar to previous reports, GUV-sized buds were abundant and close-packed on the surface of nanocellulose paper for the lipid composition consisting of pure DOPC (Fig. 2a) ^16^. The samples prepared with DOPC + 3 mol % PEG2000-DSPE did not show any apparent differences from the samples consisting of pure DOPC (Fig. 2b). Both samples had buds that appeared as 5–6 size stratified layers above the surface of the paper (Fig. 2a, b). On the glass slide, buds on the lipid film consisting of pure DOPC formed a single layer on the surface and appeared sparse compared to the buds on the surface of nanocellulose paper (Fig. 2c)^16^. The sample prepared with DOPC + 3 mol % PEG2000-DSPE on the glass slide however showed a marked difference compared to the other three samples. The lipid film formed a voluminous foam-like mesophase that extended up to 130 - 160 µm from the surface of the glass (Fig. 2d).

We highlight key areas in the image using white and red arrows and show *x - y* slices corresponding to these locations in Fig. 2e-g. The region closest to the glass consisted of a few spherical buds located between large, connected membrane septa (red arrows in Fig. 2d). The septa met at vertices of 3 to 4 septa (Fig. 2d lower panel). Further away from the surface, the mesophase had lipid dense regions, regions of separated septa, and pockets of GUV-like spherical structures. The lipid dense regions appeared web-like, with holes of low fluorescence intensity, which we interpret to be regions of continuous phase (Fig. 2e). These holes were in a lattice with hexagonal symmetry 1 µm apart (see SI Figure 4 showing zoomed images and fast Fourier transforms of the images (FFT)). The lipid dense regions appeared connected and continuous with the separated septa. In this region, the septa enclose the continuous phase forming polyhedral cells (Fig. 2f). Unlike the septa on the surface, the septa in this middle region show two clearly distinguishable membranes that appear to touch at 2 ± 0.6 µm intervals (white arrows in Fig. 2f). Further away from the surface and in regions within the mesophase, pockets of spherical GUV-like objects were evident (Fig. 2g).

Since the formation of the mesophase appears to correlate with the ultrahigh yields of GUVs, we probed the dynamics of the formation of the mesophase. We captured confocal Z-Stacks at 7 minutes, 30 minutes, and 60 minutes (Figure 3a). See Figure S5 for original images without histogram equalization. Remarkably, the mesophase was already present at 7 minutes after hydration and remains relatively unchanged at 30 minutes and 60 minutes. The GUV yield obtained by harvesting at 10 minutes was 52 ± 1 % which was statistically indistinguishable from the sample harvested at 60 minutes (*p =* 0.118). We show a histogram of the distribution of GUV diameters in Figure S5. We conclude that the GUVs originate from this foam-like mesophase and can, in principle, be obtained within 10 minutes. This mode of obtaining GUVs from a volume-spanning lamellar mesophase with the inclusion of the 3 mol % PEG2000-DSPE is distinct from surface-attached budding and merging of pure DOPC bilayers, explaining why the budding and merging model cannot explain the measured yields. The rapid formation of the mesophase is consistent with the expectation that the energy released upon hydration drives the dynamics of the lipid films on flat surfaces.^16^ Evidently, the enmeshed nanocellulose fibers of nanocellulose paper inhibits the formation of the foam-like lipid mesophase.

**Figure 3.**
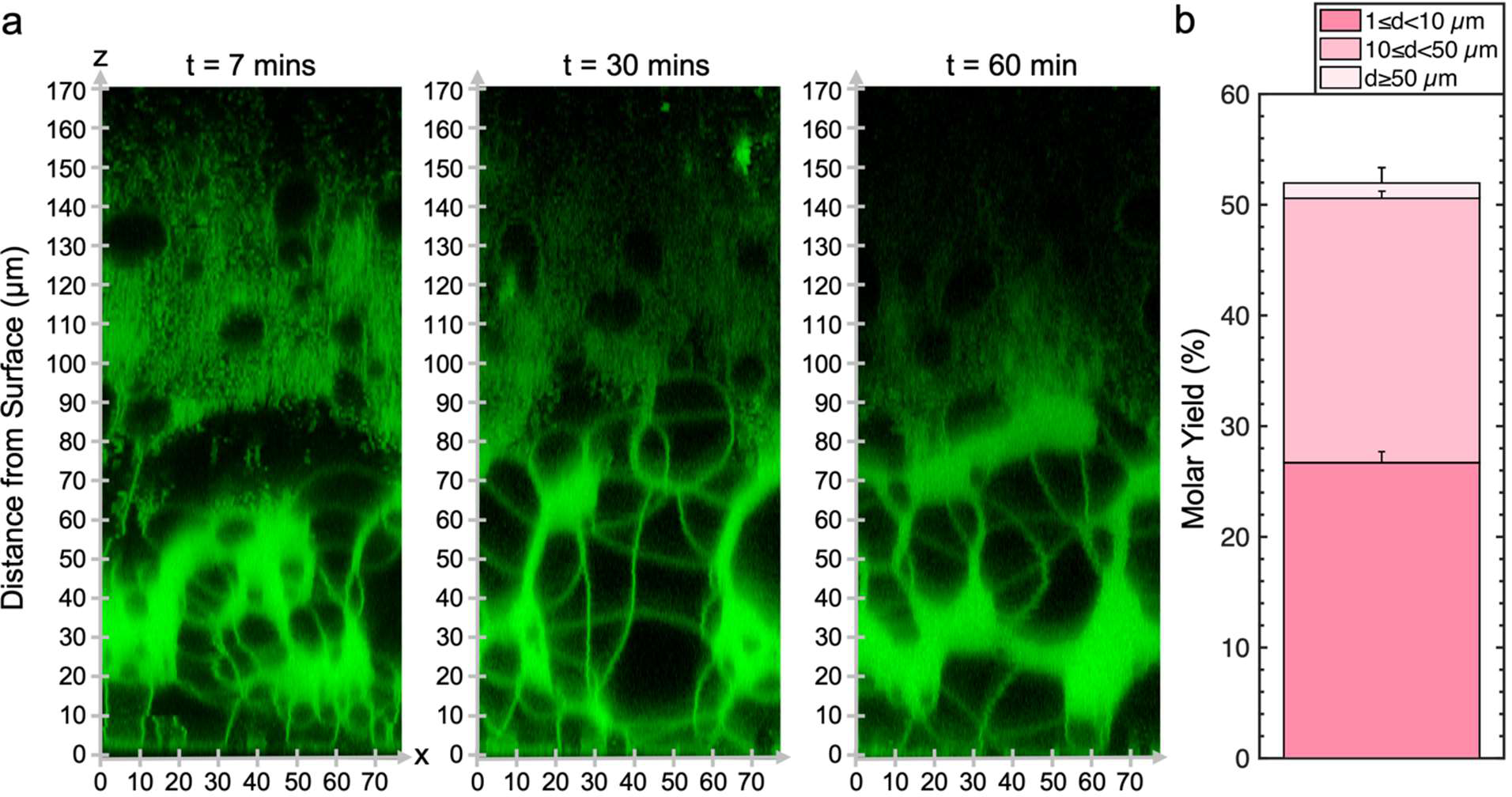
Dynamics of the lipid-dense mesophase formed from DOPC + 3 mol % PEG2000-DSPE. a) Orthogonal *x-z* planes of confocal Z-Stacks of the assembled mesophase over time. b) Molar yield of GUVs harvested after 10 minutes of incubation. The stacks show the percentage of the molar yield that is comprised of the size classifications as listed in the legend. The data is an average of N=3 independent repeats. The error bars are one standard deviation from the mean.

To understand this apparently alternative GUV formation pathway, we characterize the surface configuration of the films, the yield, and the size distributions of the harvested population of GUVs, i) with various mol % of PEG2000-DSPE, ii) with lipids that have partial molecular similarity to PEG2000-DSPE, iii) with different lipid compositions, and iv) with flat surfaces of different types.

### The configuration of the lipid film on the surfaces and the molar yields are affected by the mol % PEG2000-DSPE

We assembled GUVs using the gentle hydration method with mixtures consisting of DOPC and 0.1, 1, 5, and 10 mol % of PEG2000-DSPE. Increasing the mol % of PEG2000-DSPE increases the surface charge density of the membrane and causes the PEG2000 chain to transition from a gas of globular mushrooms to extended brushes.^34,39^ This transition is gradual and occurs when a fluid phase membrane contains 3 – 5 mol % of PEG2000-DSPE, when the distance between the PEG2000-DSPE molecules falls below twice the Flory radius, R_F_ = 3.66 nm, of PEG2000.^34,39^ High mol % of PEG2000-DSPE favors the formation of micellar phases.^18,21,39^

Confocal images show that the lipid film after 1 hour demonstrated configurations that depended on the mol % of PEG2000-DSPE (Figure 4a-c). See Figure S6 for original images without histogram equalization. In mixtures containing 0.1 and 1 mol % PEG2000-DSPE, GUV-sized buds were present on the surface. Interestingly, while there was no noticeable difference between the configuration of buds for the DOPC + 0.1 mol % PEG2000-DSPE and pure DOPC mixtures (image not shown), the DOPC +1 mol % PEG2000-DSPE mixture had 5 – 6 stacked layers of buds, similar to the configuration of the buds on the surface of nanocellulose paper (compare Fig. 4a with Fig. 2a and b). For the DOPC + 5 mol % PEG2000-DSPE mixture, a volume-spanning foam-like mesophase that was qualitatively similar to the DOPC + 3 mol % of PEG2000-DSPE mixture was present. However, this mesophase had a noticeably lower lipid density than the mesophase that formed with the DOPC+ 3 mol % PEG2000-DSPE mixture. The lower lipid density is apparent by the large fraction of continuous phase between the septa and the lack of lipid dense regions. For the DOPC + 10 mol % PEG2000-DSPE mixture, the foam-like mesophase was no longer evident. Instead, structures that appeared to be clusters of detached GUVs, the majority that have diameters < 10 µm, appeared to form. Very few buds were attached to the surface. Both the 5 mol % and 10 mol % PEG2000-DSPE had regions with small bright speckles that diffused freely (white arrows in Fig. 4b and Fig. 4c, *x-y* images in Fig. 4d and 4e). These bright speckles were not present in mixtures containing 3 mol % or lower of PEG2000-DSPE. These speckles are consistent with micelles that are known to form in mixtures with high mol % of PEG2000-DSPE^18,21^.

**Figure 4.**
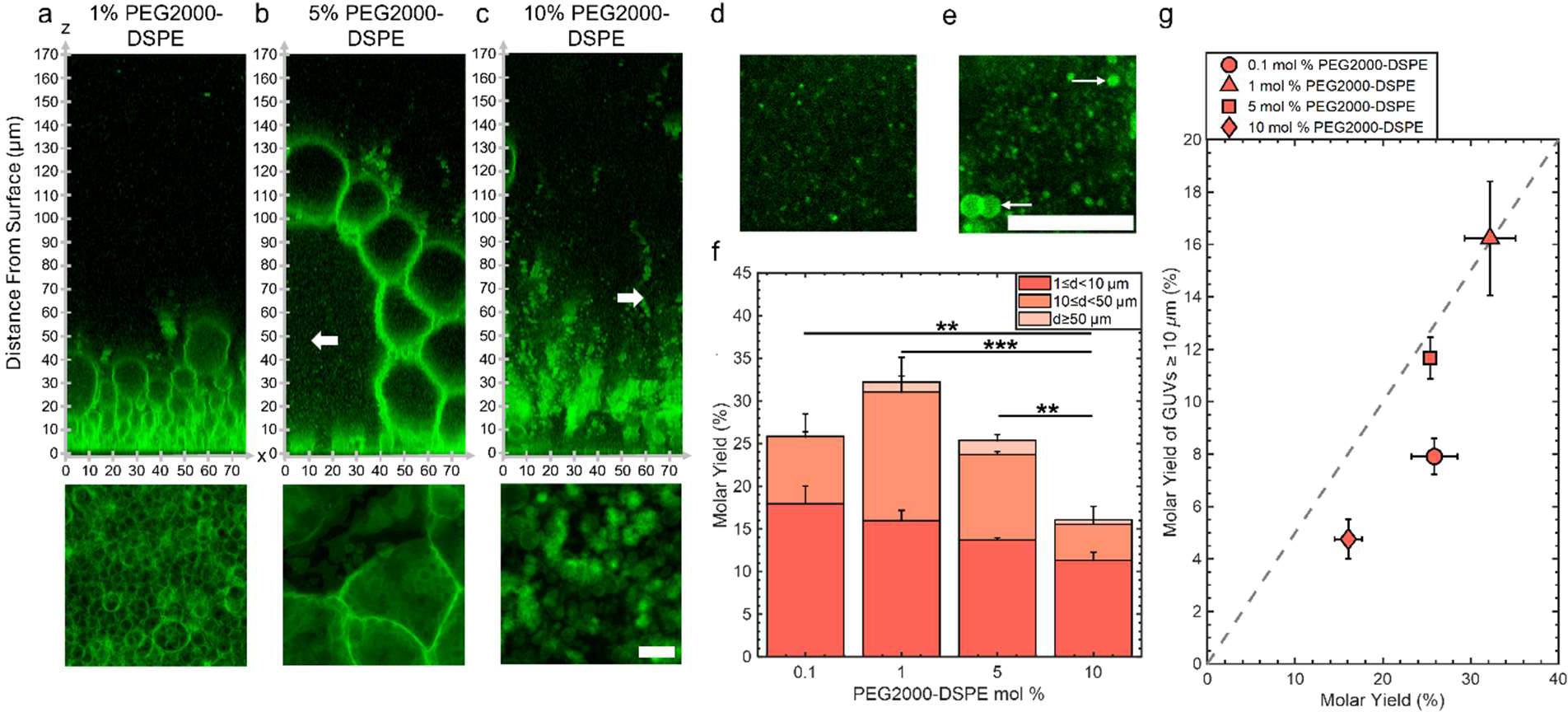
Effect of the mol % of PEG2000-DSPE on the configuration of the film and molar yields. a-c) Upper panels show orthogonal *x-z* planes of confocal Z-Stacks of the lipid films on the surface. Bottom panels are *x-y* planes consisting of summed z-projections of the first 18 µm from the surface of the substrate. a) DOPC + 1 mol % PEG2000-DSPE. b) DOPC + 5 mol % PEG2000-DSPE. c) DOPC + 10 mol % PEG2000-DSPE. Example of small micelle-like structures from the region shown by the thick white arrow in b) and c). d) 5 mol % PEG2000-DSPE. e) 10 mol % PEG2000-DSPE. The large bright objects are GUVs and MLVs (white arrows). f) Molar yields of GUVs with increasing mol % of PEG2000-DSPE. The stacks show the percentage of the molar yield that is comprised of the size classifications as listed in the legend. Each bar is an average of N=3 independent repeats. g) Scatter plot showing the molar yield from GUVs ≥ 10 µm in diameter versus the total molar yield. The grey dashed line indicates where half the lipid molecules are in GUVs with diameters ≥ 10 µm. Scale bars are 15 µm.

Figure 4f shows a plot of the molar yields obtained using the lipid mixtures. We show representative images of the harvested objects and histograms of the size distributions in Figure S7 and S8. For 0.1,1, 5, and 10 mol % of PEG2000-DSPE, the molar yield was 26 ± 3 %, 32 ± 3 %, 25 ± 1 %, and 16 ± 2 %, respectively. In addition to the DOPC + 3 mol % PEG2000-DSPE lipid mixture, only the DOPC + 1 mol % PEG2000-DSPE lipid mixture showed a yield significantly different from pure DOPC (*p* = 7.76×10^-4^). Furthermore, this yield was statistically indistinguishable from the yield of GUVs obtained using the PAPYRUS method (*p*=0.348). Figure 4g shows a plot of the yield of GUVs with *d* ≥ 10 µm versus the total molar yield of GUVs. For lipid mixtures with 0.1,1, 5, and 10 mol % of PEG2000-DSPE, the fraction of GUVs with *d* ≥ 10 µm was 31 %, 50 %, 46 %, and 30 %, respectively. Thus, the DOPC + 1 mol % PEG2000-DSPE and DOPC + 5 mol % PEG2000-DSPE lipid mixtures had a higher fraction of GUVs with *d* ≥ 10 µm compared to pure DOPC. The DOPC + 0.1 mol % PEG2000-DSPE, and DOPC + 10 mol % PEG2000-DSPE lipid mixtures had lower fraction of GUVs with *d* ≥ 10 µm compared to pure DOPC. All these mixtures had a lower fraction of GUVs with *d* ≥ 10 µm compared to pure DOPC obtained using the PAPYRUS method and compared to DOPC + 3 mol % PEG2000-DSPE obtained using the PAPYRUS and the gentle hydration methods.

To surmise, only the mixtures containing 1 and 3 mol % of PEG2000-DSPE resulted in yields significantly different from pure DOPC, and only the mixture containing 3 mol % PEG2000-DSPE had ultrahigh yields. Consistent with their similar surface configuration which consisted of a close-packed layer of size-stratified buds, the lipid mixture containing 1 mol % of PEG2000-DSPE had similar yields to the PAPYRUS method despite the flat geometry of the surface. All the other compositions, despite showing similar yields to pure DOPC, had different configurations of the film on the surfaces. The DOPC + 0.1 mol % PEG2000-DSPE lipid mixture showed similar configuration of a single sparse layer of buds to pure DOPC, the DOPC + 5 mol % PEG2000-DSPE lipid mixture had a lipid-poor foam-like mesophase and micelles, while the DOPC + 10 mol % PEG2000-DSPE lipid mixture had clusters of detached GUVs and micelles. These results show that similar yields of GUVs could arise from different configurations of the lipid film on the surfaces.

### Long-range electrostatic and short-range steric interactions are both needed for the formation of the lipid-dense foam-like mesophase and for obtaining ultrahigh yields

We investigated the molecular mechanism for the formation of the foam-like mesophase by using lipids with partial molecular similarity to PEG2000-DPSE. We prepared lipid films composed of 3 mol % DSPE, 3 mol % PEG2000-DSG, and 3 mol % DSPG. Figure 5a shows the structures of the lipids relative to PEG2000-DSPE. DSPE is the lipid portion of PEG2000-DSPE. PEG2000-DSG has a poly(ethylene glycol) chain similar to PEG2000-DSPE but no negatively charged phosphate group. DSPG has a negatively charged phosphate group similar to the negatively charged phosphate group of PEG2000-DSPE but no poly(ethylene glycol) chain. Confocal images show that the films form a layer of surface-attached buds for all three lipid mixtures. There is no foam-like mesophase (Fig. 5b, see Figure S9 for images without histogram equalization). Furthermore, we find that none of the lipid mixtures resulted in GUV yields significantly different than pure DOPC, albeit the distributions in sizes were different (Figure 6a). We find that the addition of DSPE resulted in a yield of 16 ± 0.5 %. The molar yield for PEG2000-DSG was not significantly different from DSPE at 17 ± 2.5 %. Upon the inclusion of DSPG, the molar yield increased significantly in comparison to both DSPE and PEG2000-DSG (both *p* < 0.05) to 26 ± 2 %. The fraction of GUVs with *d* ≥ 10 µm for DSPE, DSPG and PEG2000-DSG was 25 %, 48 %, and 35 % respectively (Fig. 6b).

**Figure 5.**
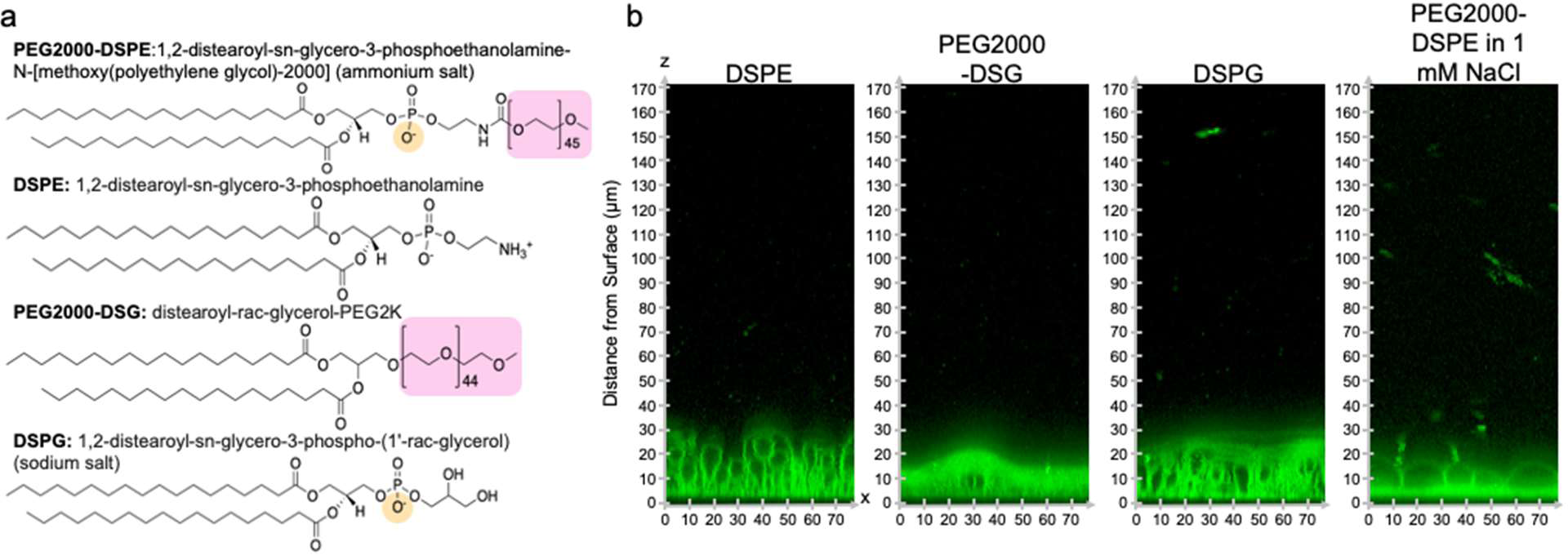
Chemical structures of lipids and configuration of the films on the surface. a) Chemical structures of lipids used. The PEG chain is highlighted in pink and the negative charge is highlighted in yellow. b) Orthogonal *x-z* planes of confocal Z-Stacks of the lipid films on the surface after 1 hour of hydration.

**Figure 6.**
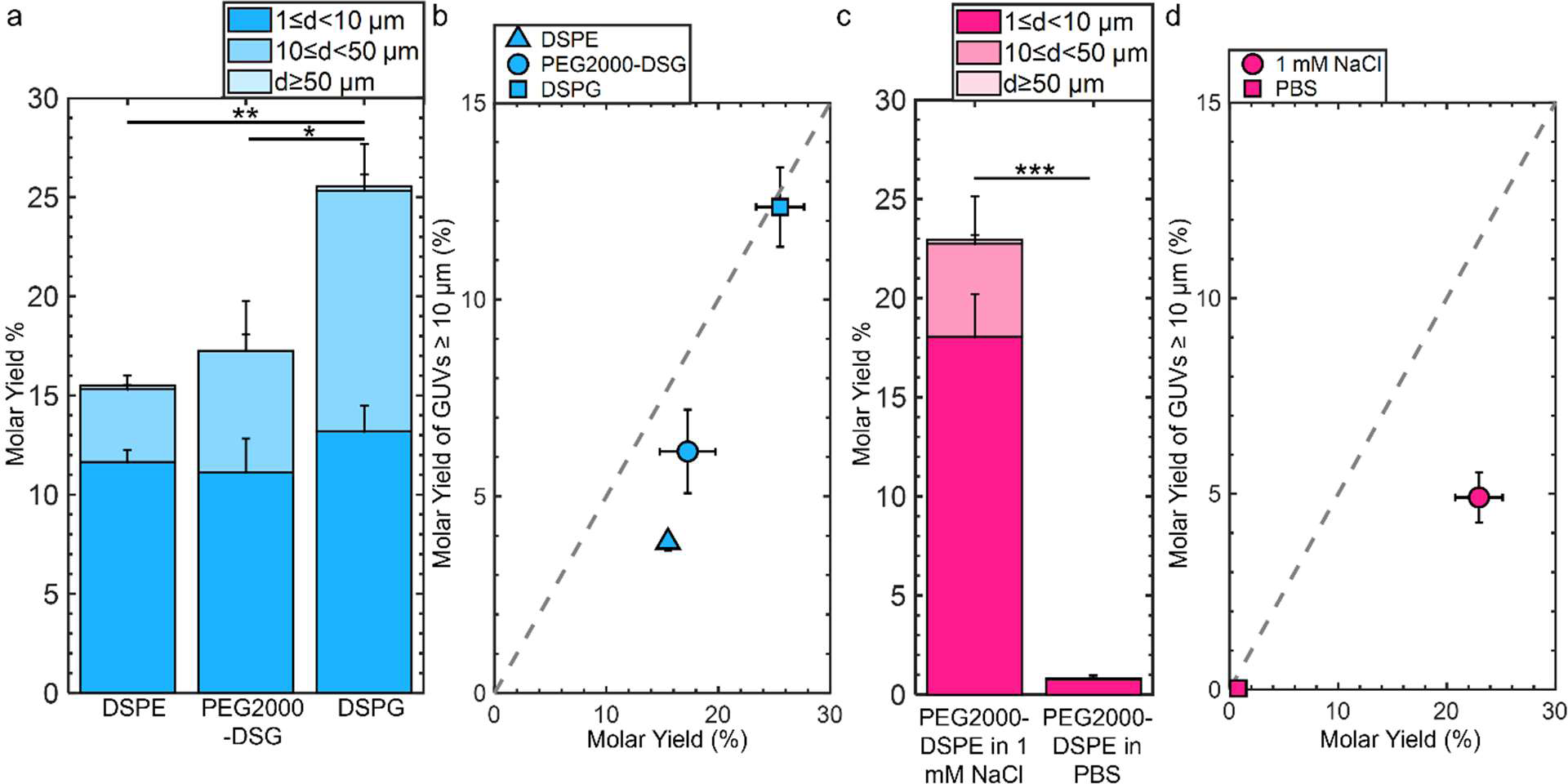
Molar yields of GUVs assembled with lipids with partial molecular similarity to PEG2000-DSPE. a) Stacked bar plots showing molar yields from the gentle hydration method with DOPC + 3 mol % DSPE, DOPC + 3 mol % PEG2000-DSG, and DOPC + 3 mol %. b) Scatter plot showing the molar yield from GUVs ≥ 10 µm in diameter versus the total molar yield for DOPC + 3 mol DSPE, DOPC + 3 mol PEG2000-DSG and DOPC + 3 mol DSPG. c) Stacked bar plots showing molar yields from the gentle hydration method with DOPC + 3 mol % PEG2000-DSPE in 100 mM sucrose + 1 mM NaCl and 100 mM sucrose + PBS. d) Scatter plot showing the molar yield from GUVs ≥ 10 µm in diameter versus the total molar yield. The grey dashed line indicates where half the lipid molecules are in GUVs with diameters ≥ 10 µm. The stacks show the percentage of the molar yield that is comprised of the size classifications as listed in the legend. Each bar is an average of N=3 independent repeats. The error bars are one standard deviation from the mean. Statistical significance was determined using a Student’s *t*-test. *: p < 0.05, **: p < 0.01, ***: p < 0.001, n.s.: not significant.

Since PEG2000-DSG had no effect on the yield of GUVs, we reasoned that the formation of the foam-like mesophase and the increase in yield requires the charge group in addition to the poly(ethylene glycol) chain. he range of the electrostatic interaction is highly sensitive to the ionic strength of the solution. The Debye screening length provides an estimate of the range of electrostatic interaction.^56^ At the Debye length, the electrostatic potential decays by 1/*e*. Thus, screening lengths with smaller magnitudes reflect a shorter range at which electrostatic interactions are felt. To test the relative importance of the range of the electrostatic interaction for forming the foam-like mesophase, we hydrated films of DOPC + 3 mol % PEG2000-DSPE in 100 mM sucrose + 1 mM NaCl and in 100 mM sucrose + phosphate buffered saline (PBS) (composition: 137 mM NaCl, 2.7 mM KCl, 8 mM Na_2_HPO_4_, and 2 mM KH_2_PO_4_). The Debye screening lengths are 170 nm, 9.6 nm, and 0.75 nm in 100 mM sucrose, 100 mM sucrose + 1 mM of NaCl, and 100 mM sucrose + PBS, respectively. Thus, in these solutions, the screening lengths are ∼ 37×, 2×, and 0.2× the Flory radius of PEG2000-DSPE. The yield of GUVs dropped to 23 ± 2% in the solution with 1 mM NaCl (Fig. 6c). Similar to our previously reported result^12^, in PBS, the yield of GUVs was 0.8 ± 0.2 %. Upon the inclusion of salt, the fraction of molar yield from GUVs with *d* ≥ 10 µm reduced to 21 % in the solution containing 1 mM NaCl and to 4 % in PBS (Fig. 6d). See Figure S10-S11 for representative images of the harvested objects and histograms of the sizes.

We conclude that the formation of the foam-like mesophase is unique to PEG2000-DSPE and requires both the short-range steric repulsion of the poly(ethylene glycol) chain and the long-range repulsion of the phosphate group. In molecularly similar lipids lacking one of these interactions, or when the range of electrostatic repulsion is reduced by screening in salty solutions, the mesophase does not form. Instead, GUVs assemble through budding and merging on the surface.

### Addition of PEG2000-DSPE increases yields in complex lipid mixtures and on stainless steel substrates

We determine the generality of the strategy of incorporating 3 mol % PEG2000-DSPE for increasing GUV yields using the gentle hydration method. We use a ternary mixture consisting of the unsaturated lipid DOPC, the saturated lipid DPPC, and cholesterol as our model lipid composition. This lipid composition is a canonical mixture used in lipid biophysics for studying lateral phase separation and membrane organization^46–48^. Without PEG2000-DSPE, the molar yield was 21 ± 1 % (Fig. 7a). Incorporation of 3 mol % PEG2000-DSPE results in a 1.7× increase in the molar yield to 36 ± 3 %. We find that the size distribution additionally changes to include a large fraction of GUVs with *d* ≥ 10 µm (Fig. 7b). Without PEG2000-DSPE, 25 % of the molar yield was from GUVs with *d* ≥ 10 µm. Upon inclusion of PEG2000-DSPE, the fraction of the molar yield from GUVs with *d* ≥ 10 µm increased to 35 %.

**Figure 7.**
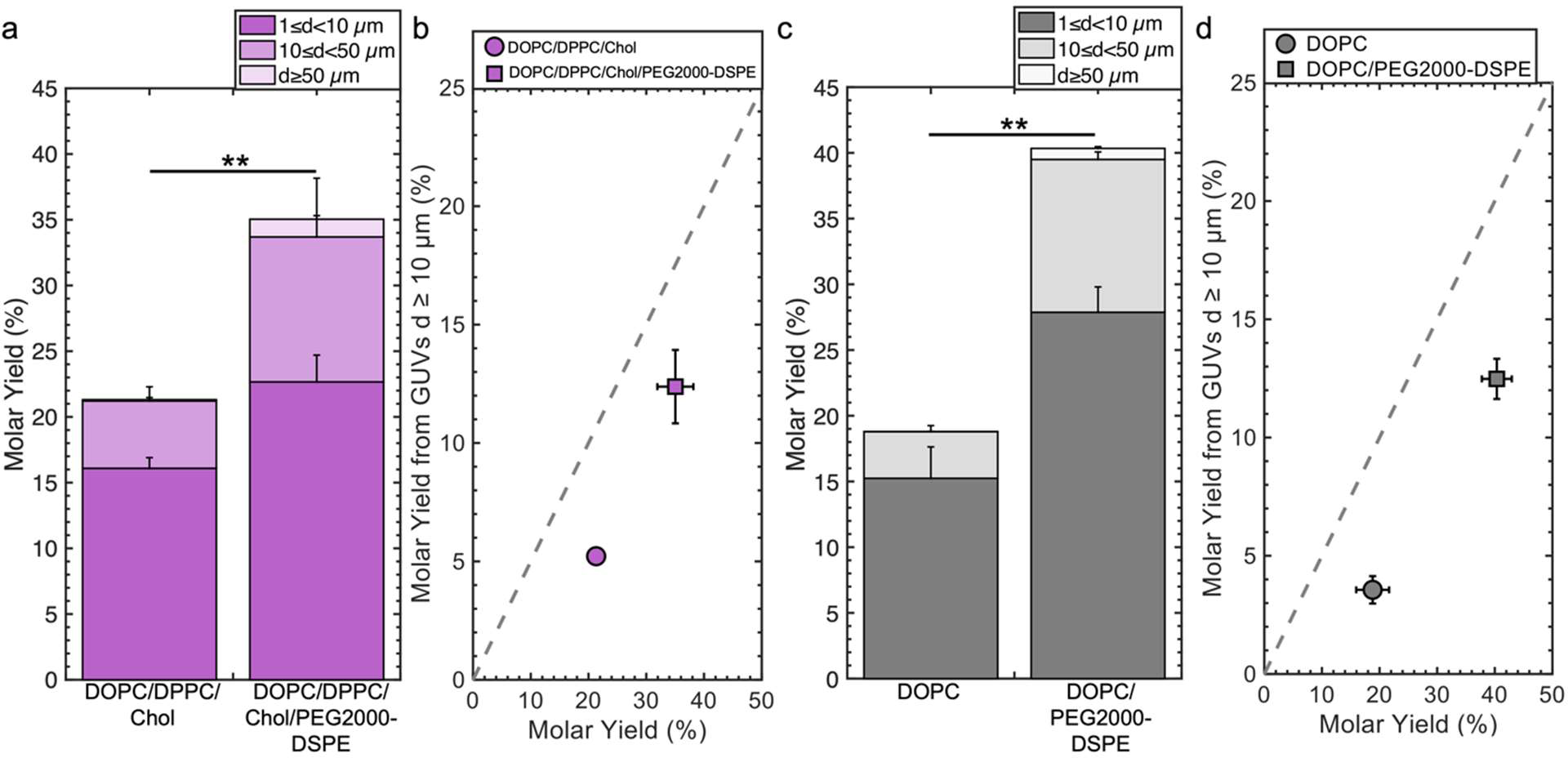
Generality of increasing yields by adding 3 mol % of PEG2000-DSPE. a) Molar yield of GUVs composed of DPPC/DPPC/Cholesterol and DOPC/DPPC/Cholesterol + 3 mol % PEG2000-DSPE. b) Scatter plot showing the molar yield from GUVs ≥ 10 µm in diameter versus the total molar yield for the samples in a). c) Molar yield of GUVs composed of pure DOPC and DOPC + 3 mol % PEG2000-DSPE obtained using the gentle hydration method on stainless steel sheets. d) Scatter plot showing the molar yield from GUVs ≥ 10 µm in diameter versus the total molar yield for the samples in c). The stacks show the percentage of the molar yield that is comprised of the size classifications as listed in the legend. Each bar is an average of N=3 independent repeats. The error bars are one standard deviation from the mean. The grey dashed line indicates where half the lipid molecules are in GUVs with diameters ≥ 10 µm. Statistical significance was determined using a Student’s *t*-test. *: p < 0.05, **: p < 0.01, ***: p < 0.001, n.s.: not significant.

We determined the generality of using other planar substrates to assemble GUVs by using stainless steel sheets. Stainless steel is a durable material used to fabricate sterilizable tools for many biomedical applications^57^. The gentle hydration method using stainless steel sheets with pure DOPC results in a yield of 19 ± 3 % (Fig. 7c). This yield is similar to the yield obtained using glass slides as a substrate. With 3 mol % PEG2000-DSPE in the lipid mixture, the yield increases to 40 ± 3 %. The yield thus is double the yield without PEG2000-DSPE, but lower than the yield obtained using glass slides as a substrate. 19 % of the GUVs had *d* ≥ 10 µm without PEG2000-DSPE whereas 31 % had *d ≥* 10 µm upon the inclusion of PEG2000-DSPE (Fig. 7d). We show histograms of the sizes in Figure S12. Thus, adding PEG2000-DSPE to lipid mixtures could be a general strategy to increase the yields obtained using the gentle hydration method on planar surfaces in low salt solutions.

### Shear induced fragmentation of the foam-like mesophase contributes to the formation of ultrahigh yields of GUVs in mixtures containing 3 mol % PEG2000-DSPE

We show in Figure 8 schematics of the proposed mechanism of the assembly of GUVs in the presence of PEG2000-DSPE. From 0 to 1 mol % PEG2000-DSPE, GUV-sized buds assemble through budding and merging on the surface (Fig. 8a, b). We propose that concentrations of 0.1 mol % and lower do not change physical parameters of the bilayer enough to result in statistically significant changes in the yield compared to using pure DOPC (Figure 8a). At 1 mol % of PEG2000-DSPE, the close-packed size-stratified layer of buds indicates the formation of many buds (Fig. 8b).

**Figure 8.**
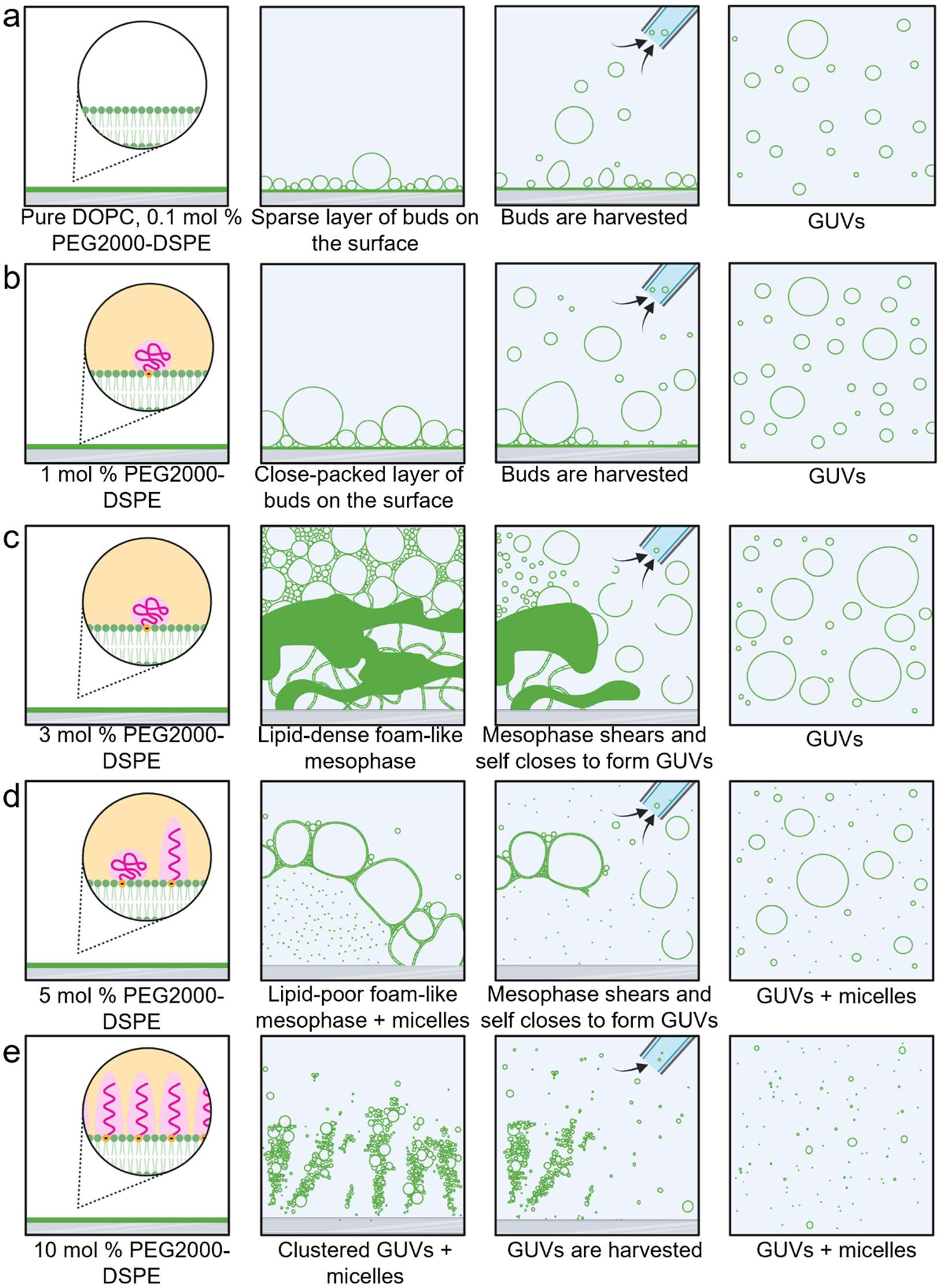
Schematic of pathways of assembly of GUVs in lipid mixtures containing PEG2000-DPSE. a) pure DOPC and DOPC + 0.1 mol % PEG2000-DSPE. b) DOPC + 1 mol % PEG2000-DSPE. c) DOPC + 3 mol % PEG2000-DSPE. d) DOPC + 5 mol % PEG2000-DSPE. e) DOPC + 10 mol % PEG2000-DSPE.

At the higher mol fraction of 3 mol % PEG2000-DSPE, we propose that the high repulsive forces contributed by the long-ranged electrostatic interactions and short-ranged steric interactions causes the rapid expansion of the film to form a lipid-dense foam-like mesophase (Fig 8c). GUVs assemble due to shear-induced breakup and closure of the membranes in the mesophase when the pipette is used for harvesting. The median size of GUVs, 3.9 ± 0.1 µm, is smaller than the median size of the polyhedral cells in the mesophase, 25 ± 10 µm. This result is consistent with the mesophase fragmenting and self-closing to form GUVs upon application of fluid shear.

At 5 mol % of PEG2000-DSPE (Fig 8d), micellization reduces the lipid density in lamellar membranes in favor of micelles, and at 10 mol % (Fig 8e), the mesophase appears to lose integrity. We propose that at this high mol % of PEG2000-DSPE, the lipids are mainly in micelles and only small clusters of GUVs remain on the surface.

Having a substrate with flat geometry is essential for the formation of the foam-like mesophase. The cylindrical geometry and the entangled fibers of nanocellulose paper appear to prevent the formation of the mesophase. Instead, buds evolve on the surface due to budding and merging even when 3 mol % or PEG2000-DSPE is present in the membrane.

### Conclusion

This work shows that incorporation of PEG2000-DSPE results in complex but measurable effects on the GUV yield and configuration of the film on the surface. The complex effects belie simplistic correlation to changes in a single physical parameter in the membrane. At 1 mol % of PEG2000-DSPE, there are multiple layers of GUV-sized buds on the surface, while at 3 mol % PEG2000-DSPE, a foam-like lipid dense mesophase forms that is broken up during harvesting. Breakup of this lipid dense mesophase is a pathway of assembly that can result in ultrahigh yields of GUVs. Incorporating 3 mol % of PEG2000-DSPE in a phase separating mix and on stainless steel substrates is effective in doubling the yields of GUVs compared to mixtures without PEG2000-DSPE. This effect appears to be unique to PEG2000-DSPE and does not occur in lipids with partial molecular similarity such as those that only have a charged phosphate headgroup or only a PEG2000 chain. Furthermore, when range of the electrostatic charge of the phosphate headgroup on PEG2000-DSPE is screened, the yield of GUVs falls. The yield of GUVs becomes negligible in solutions with physiological ionic strengths.

From a mechanistic perspective, this work reports the discovery of an alternate pathway for forming GUVs via mesophase break up in membranes containing 3 mol % PEG2000-DSPE. This pathway is distinct from budding and merging on surfaces.^16^ Thus, two different pathways that depend on the membrane composition and the surface geometry result in GUVs.

Looking forward, we anticipate other factors such as the surface concentration of lipid and the composition of the membrane could favor the formation of GUVs via one pathway over the other. The framework that couples quantitative measurements of yield with direct high-resolution visualization of the lipid film that we use here will be useful for understanding other systems. Furthermore, PEG2000-DSPE in the range of 1 to 3 mol % is used widely in biomedical applications^22,23,25^ and surfaces such as stainless steel are used in bioreactor chambers.^57^ Our results show that in addition to conferring essential properties such as “stealth” capabilities and increased circulation lifetime, addition of 1 to 3 mol % of PEG2000-DSPE into lipid mixtures provides a route to obtaining high and ultrahigh yields of GUVs via the gentle hydration method using myriad surfaces for potential uses in biomedicine.

## ASSOCIATED CONTENT

### Supporting Information

The Supporting Information is available at DOI:

Supporting text describing:

Figures showing: Representative images of harvested GUVs, orthogonal Z-stack reconstructions without histogram equalization, and histograms of GUV size distributions.

Tables showing: F and *p*-values of statistical tests.

Figure S1-S12 and Table S1-S6 (PDF).

### Author contributions

A.B.S. conceived and directed the study. A.C. performed experiments and analyzed the data. A.B.S. and A.C. interpreted the data. A.B.S. proposed the shear induced fragmentation mechanism. A.C. prepared the figures. A.B.S and A.C. wrote the manuscript. All authors have given approval to the final version of the manuscript.

## Acknowledgements

This work was funded by the National Science Foundation through NSF CAREER DMR-1848573. The data in this work was collected, in part, with a confocal microscope acquired through the National Science Foundation MRI Award Number DMR-1625733.

## Supporting Information

### Calculation of Debye screening length

We follow reference^1^ to calculate the Debye screening length. The range of the electrostatic repulsion depends on the Debye length, 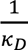, where *K_D_* is:

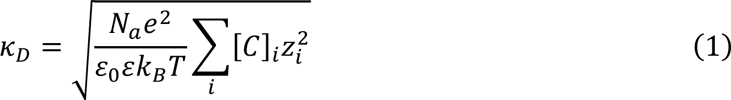

In this equation, *N_A_* is Avogadro’s number, *e* is the elementary charge, ε_0_ is the permittivity of free space, ε is the dielectric constant, *k_B_* is the Boltzmann constant, *T* is the absolute temperature, [*C*] is the concentration of ionic species *i*, and *j* is the charge of ionic species *i*. In unbuffered ultrapure water with dissolved nonionic solutes in equilibrium with atmosphere, the Debye length is ∼ 170 nm while in 1 mM NaCl it is 9.6 nm. In 1× PBS, the Debye length is 0.75 nm.

## Supporting Figures

**Figure S1.**
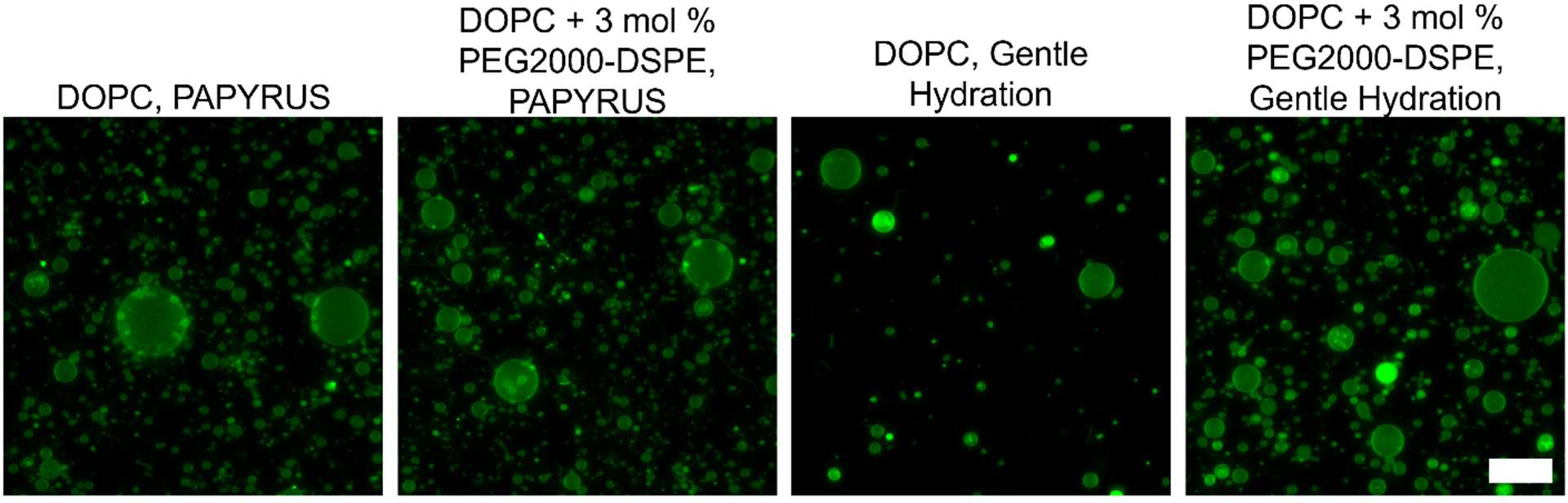
Representative images of harvested GUVs from samples shown in Figure 1. The scale bar is 50 µm.

**Figure S2.**
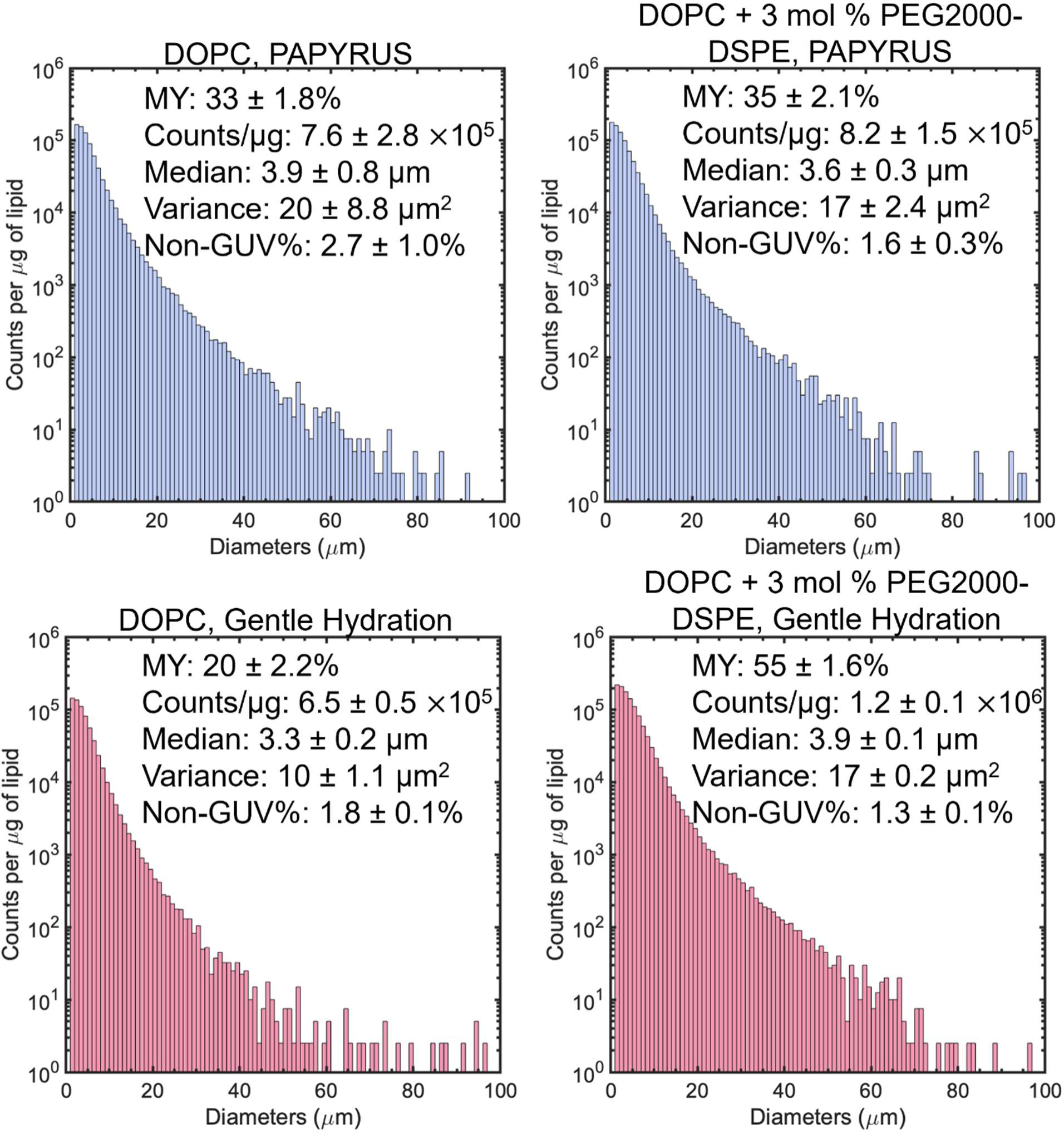
Histograms of GUV diameters of the samples shown in Figure 1. Each histogram is the average of N=3 independent repeats. Note the logarithmic scale on the *y*-axis. Bin widths are 1 µm. The molar yield (MY), total counts per µg of lipid (counts/ µg), median diameter, non-GUV % and variance are reported in the text inserts.

**Figure S3.**
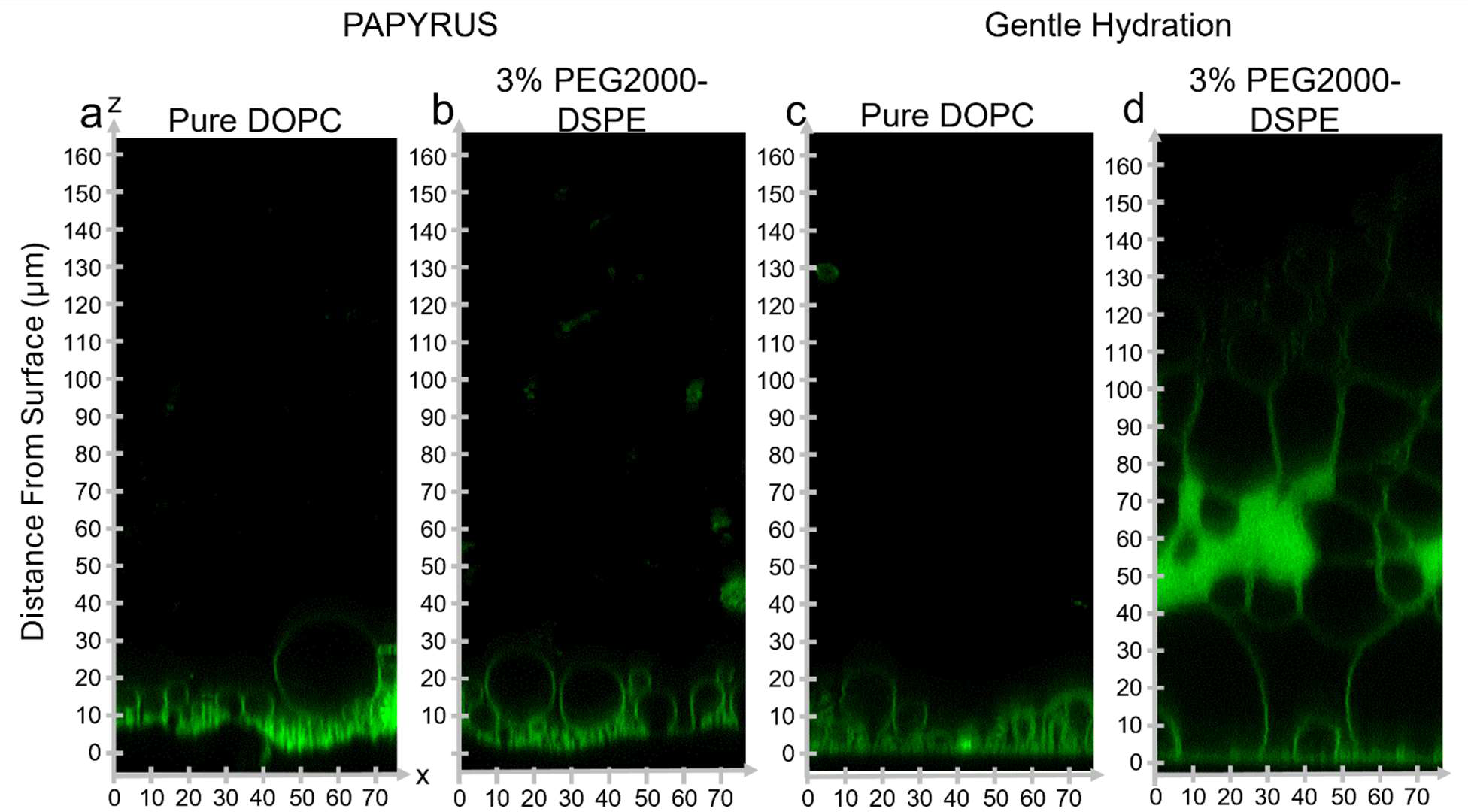
Orthogonal *x-z* images shown in Figure 2 without histogram equalization.

**Figure S4.**
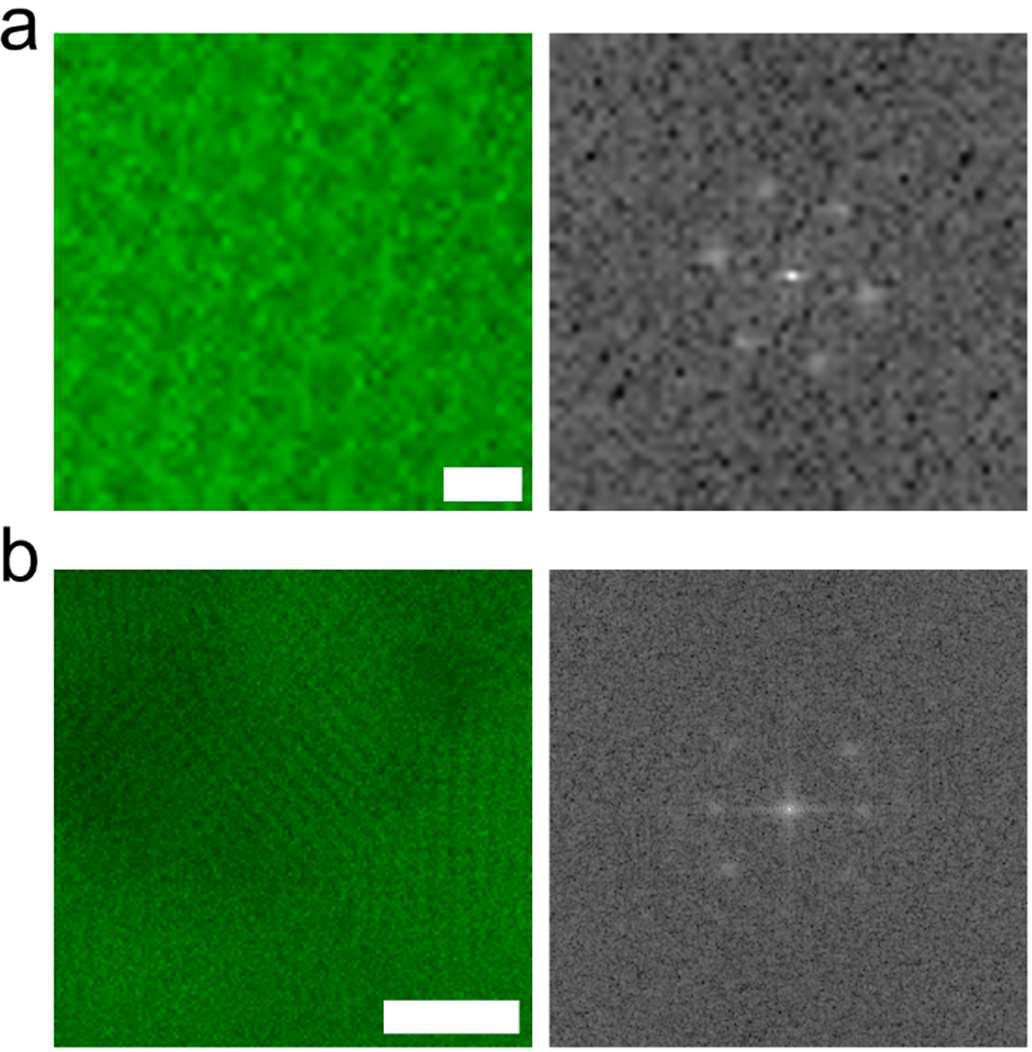
Zoomed images of the lipid dense region (left) and its corresponding fast Fourier transform, FFT (right). Scale bars are a) 1 µm and b) 5 µm.

**Figure S5.**
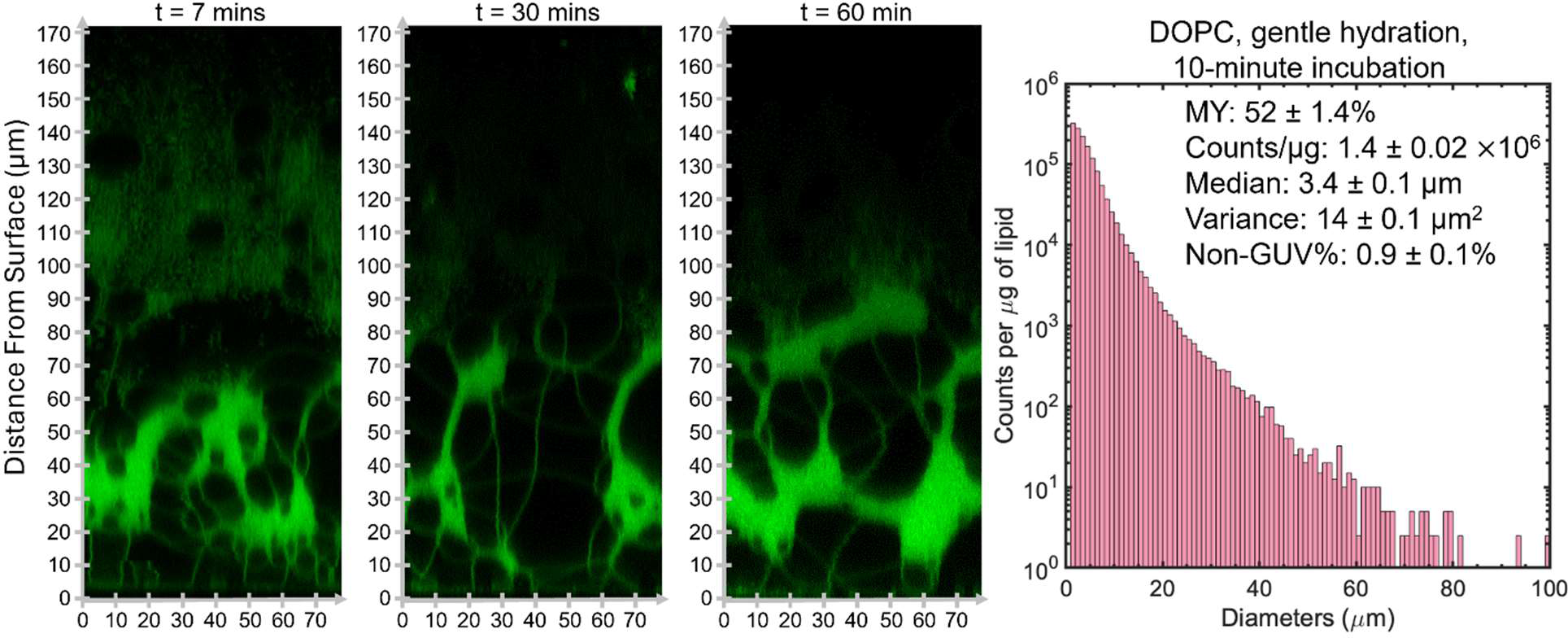
Orthogonal *x-z* images without histogram equalization shown in Figure 3 and histogram of GUV diameters. The histogram is the average of N=3 independent repeats. Note the logarithmic scale on the *y*-axis. Bin widths are 1 µm. The molar yield (MY), total counts per µg of lipid (counts/ µg), median diameter, non-GUV% and variance are shown in the text insert on the plots.

**Figure S6.**
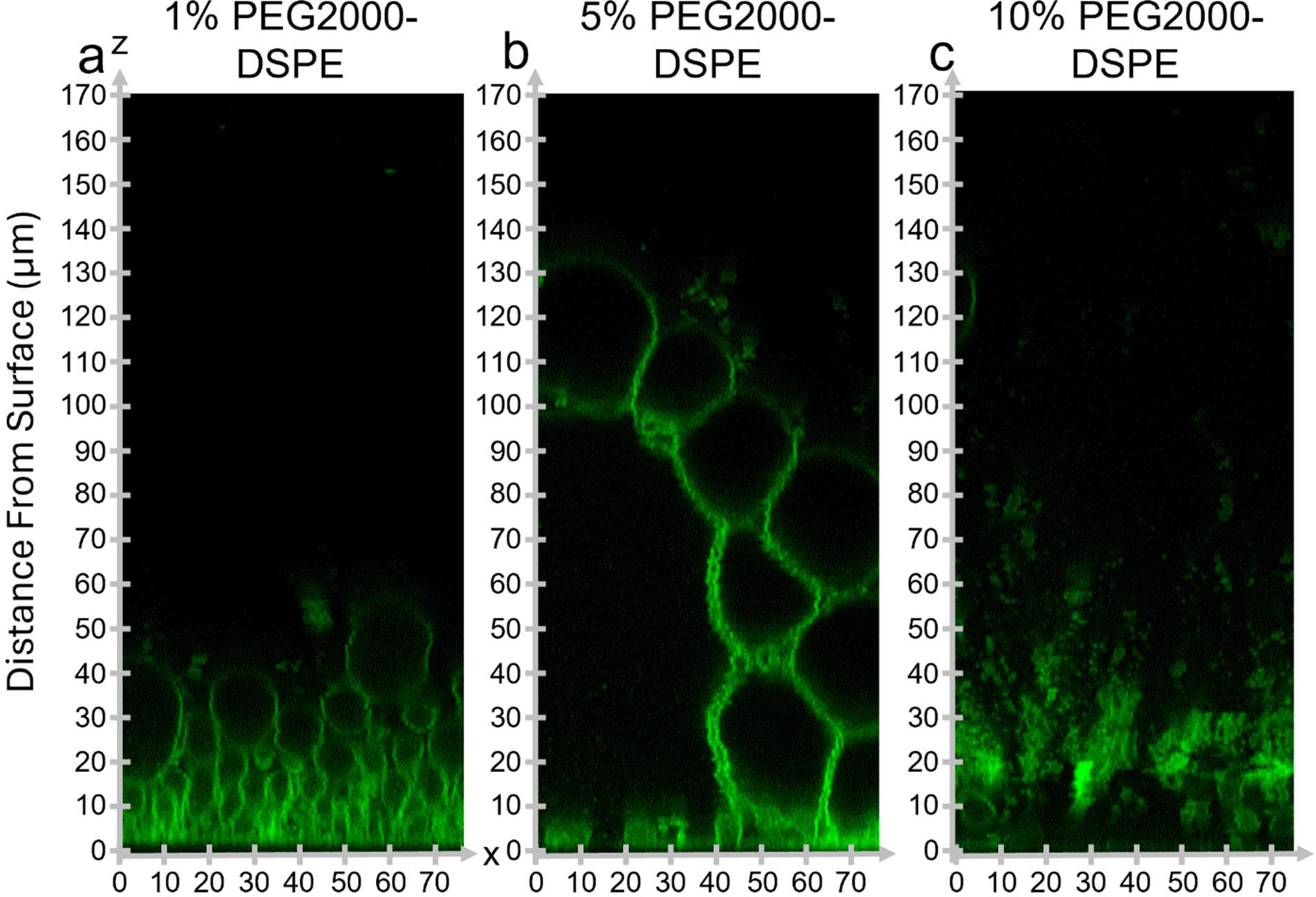
Orthogonal *x-z* images without histogram equalization shown in Figure 4.

**Figure S7.**
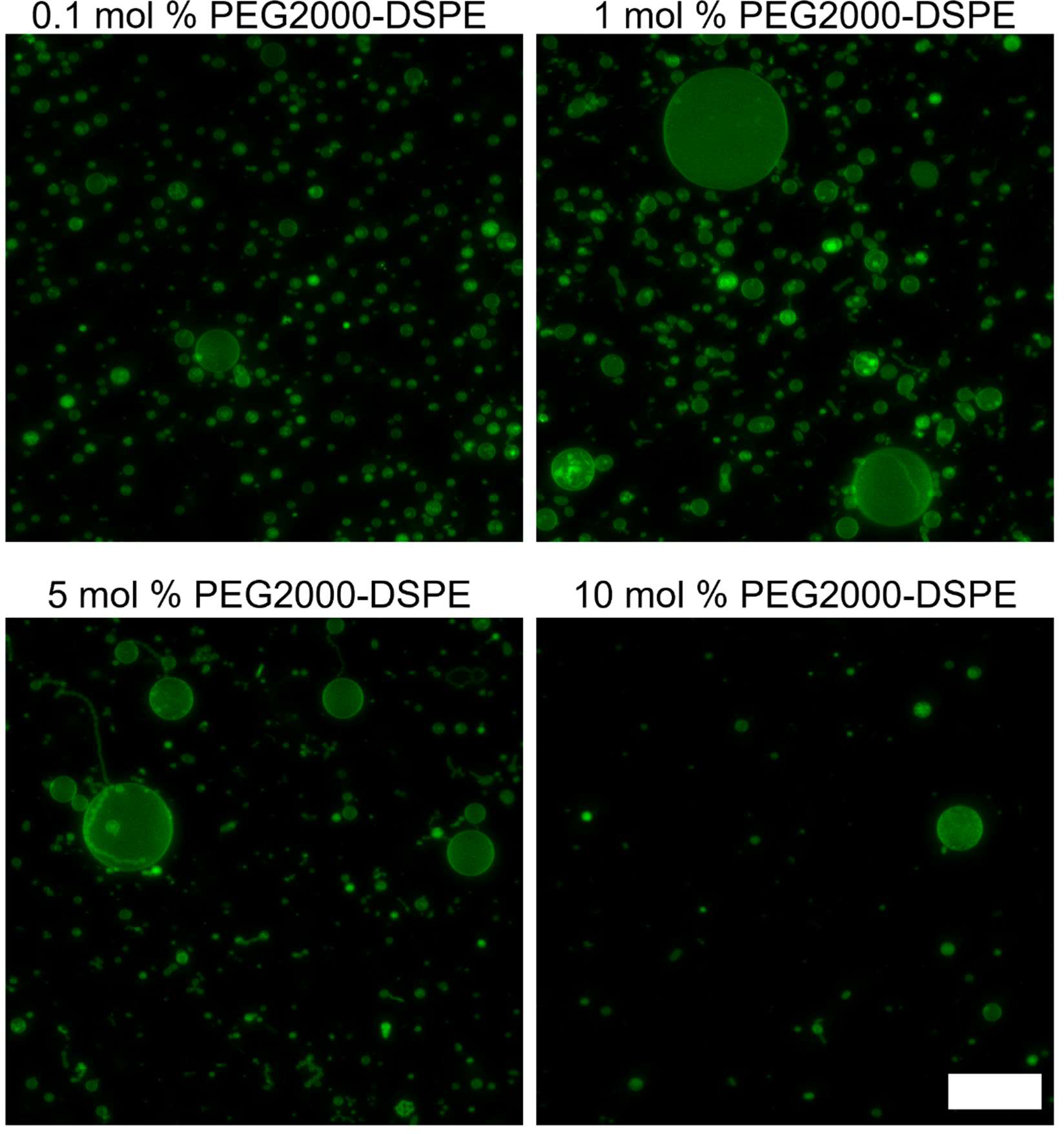
Representative images of harvested GUVs from the samples shown in Figure 4f. The scale bar is 50 µm.

**Figure S8.**
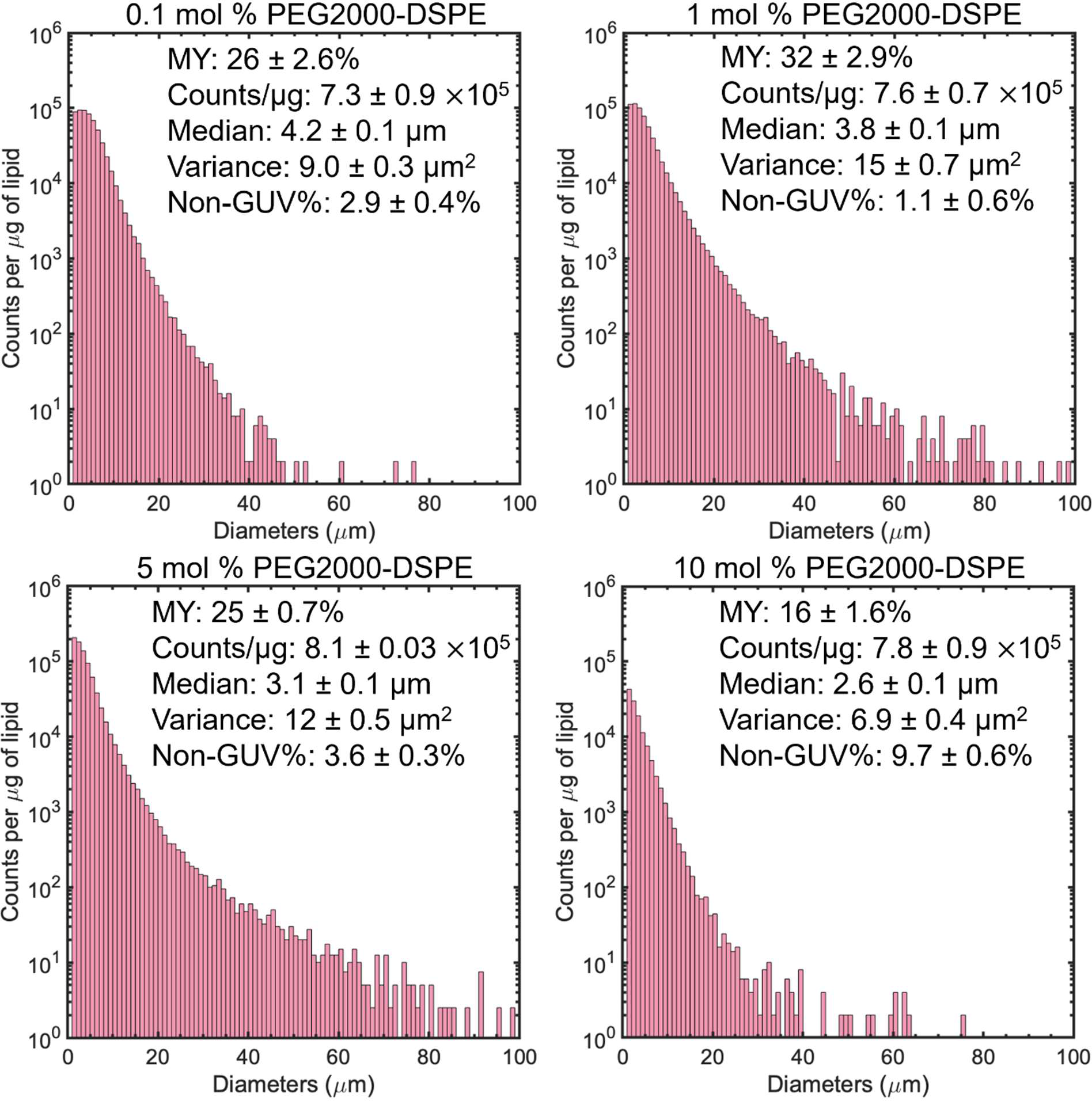
Histograms of GUV diameters of samples shown in Figure 4f. Each histogram is the average of N=3 independent repeats per sample. Note the logarithmic scale on the y-axis. Bin widths are 1 µm. The molar yield (MY), total counts per µg of lipid (counts/ µg), median diameter, non-GUV% and variance are shown in the text insert on the plots.

**Figure S9.**
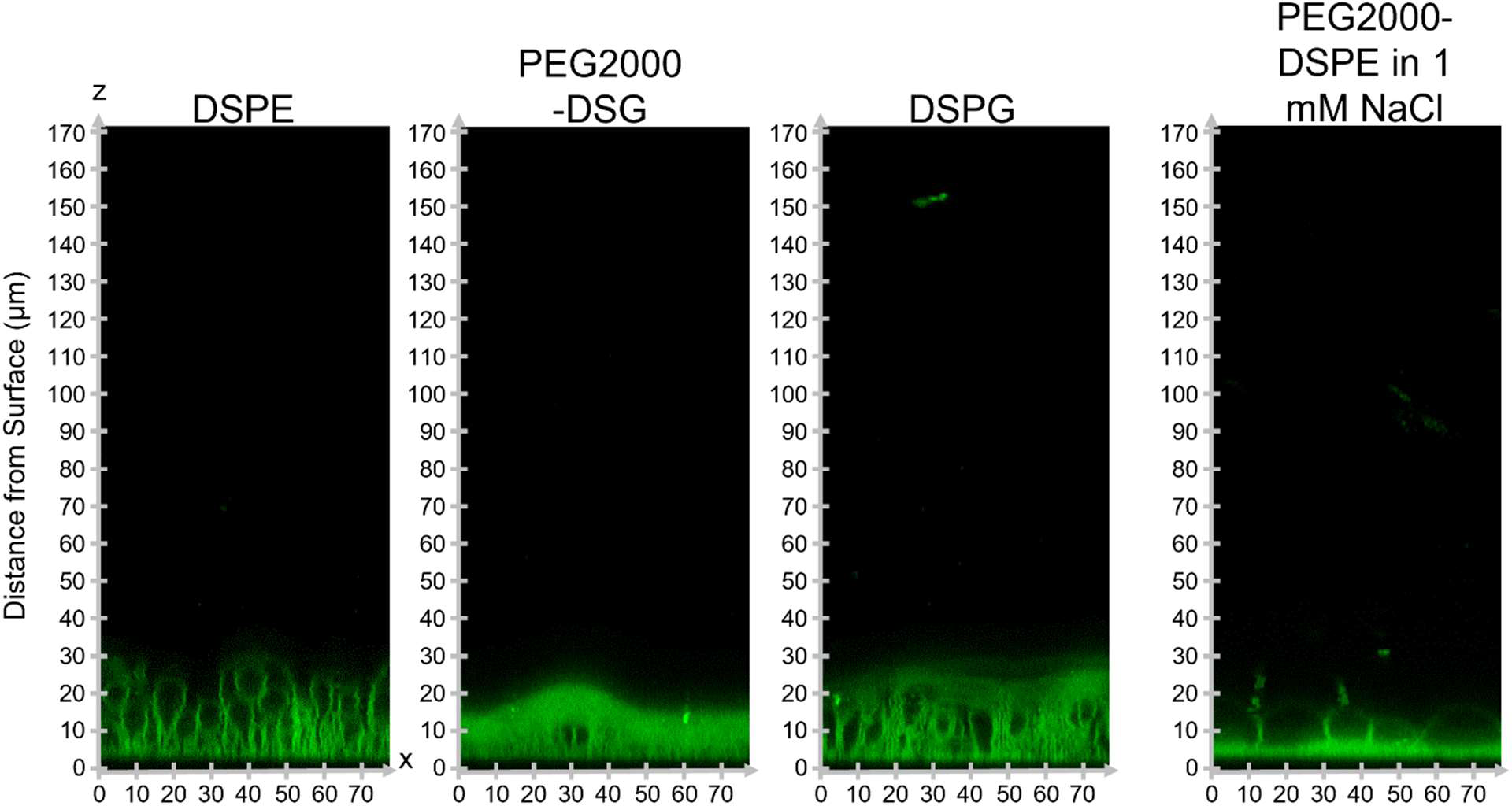
Orthogonal *x-z* images without histogram equalization shown in Figure 5.

**Figure S10.**
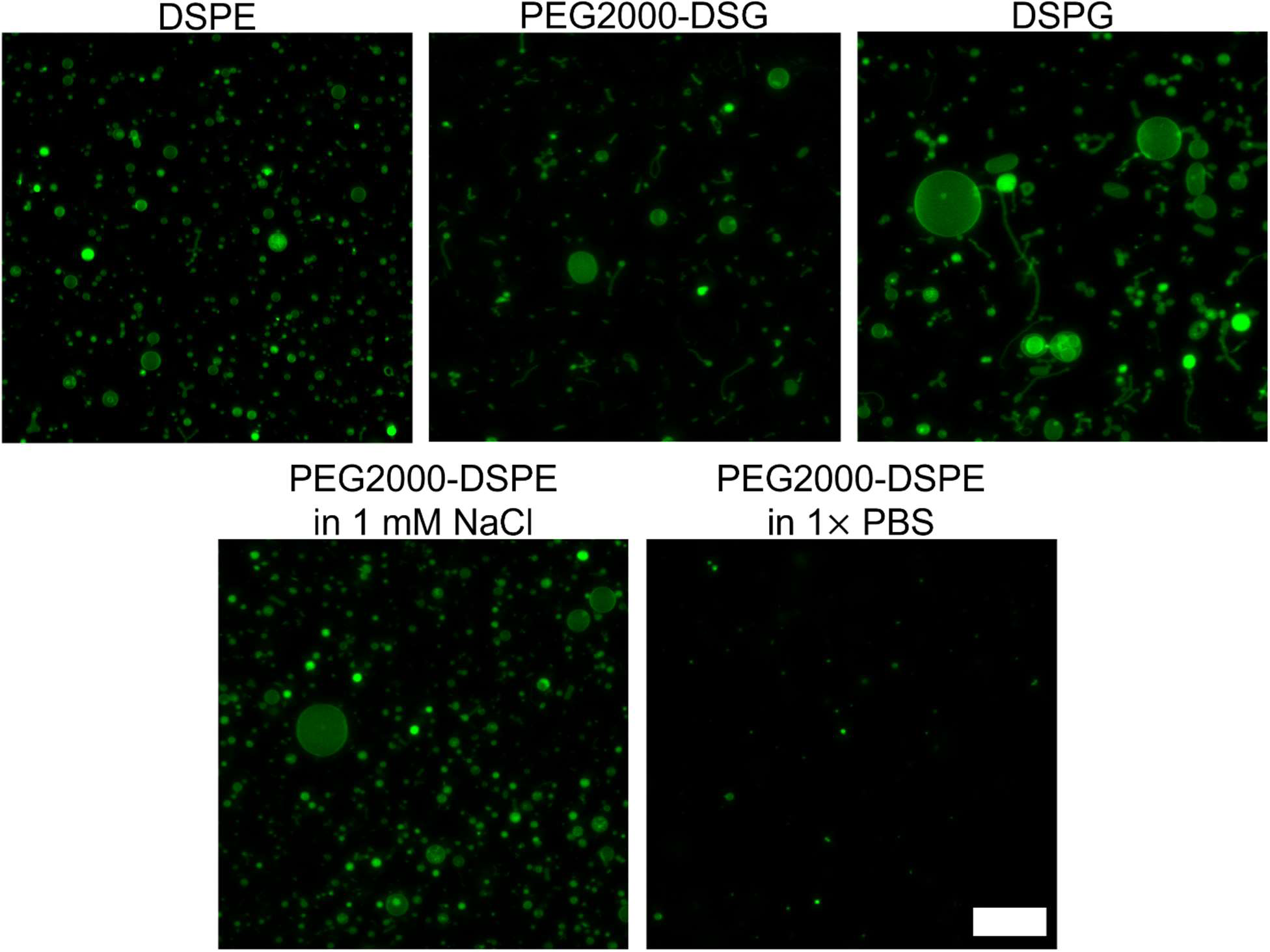
Representative harvested images from the samples imaged for Figure 6. Scale bar is 50 µm.

**Figure S11.**
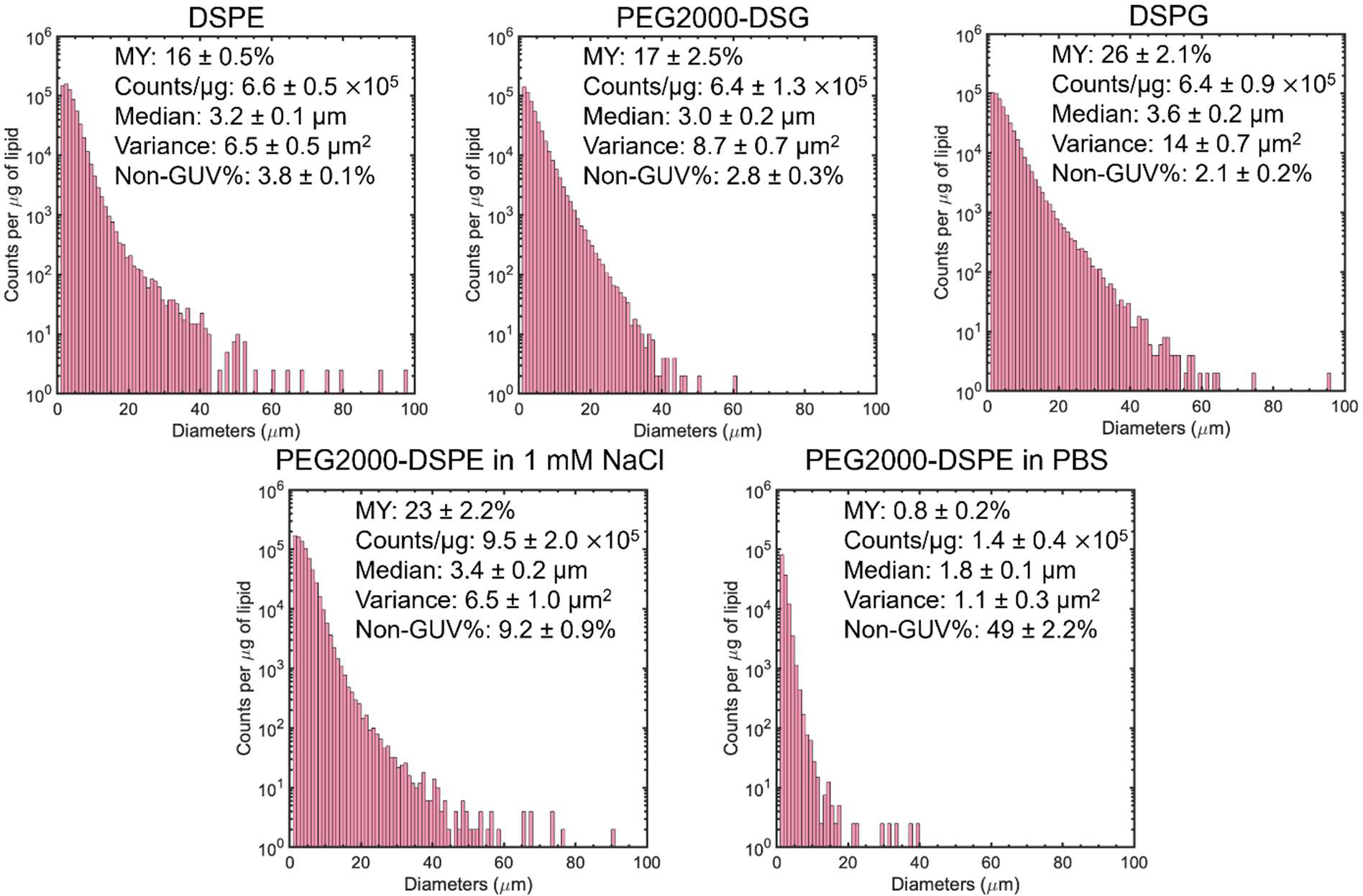
Histograms of GUV diameters of samples shown in Figure 6. Each histogram is the average of N=3 independent repeats per sample. Note the logarithmic scale on the y-axis. Bin widths are 1 µm. The molar yield (MY), total counts per µg of lipid (counts/ µg), median diameter, non-GUV% and variance are shown in the text insert on the plots.

**Figure S12.**
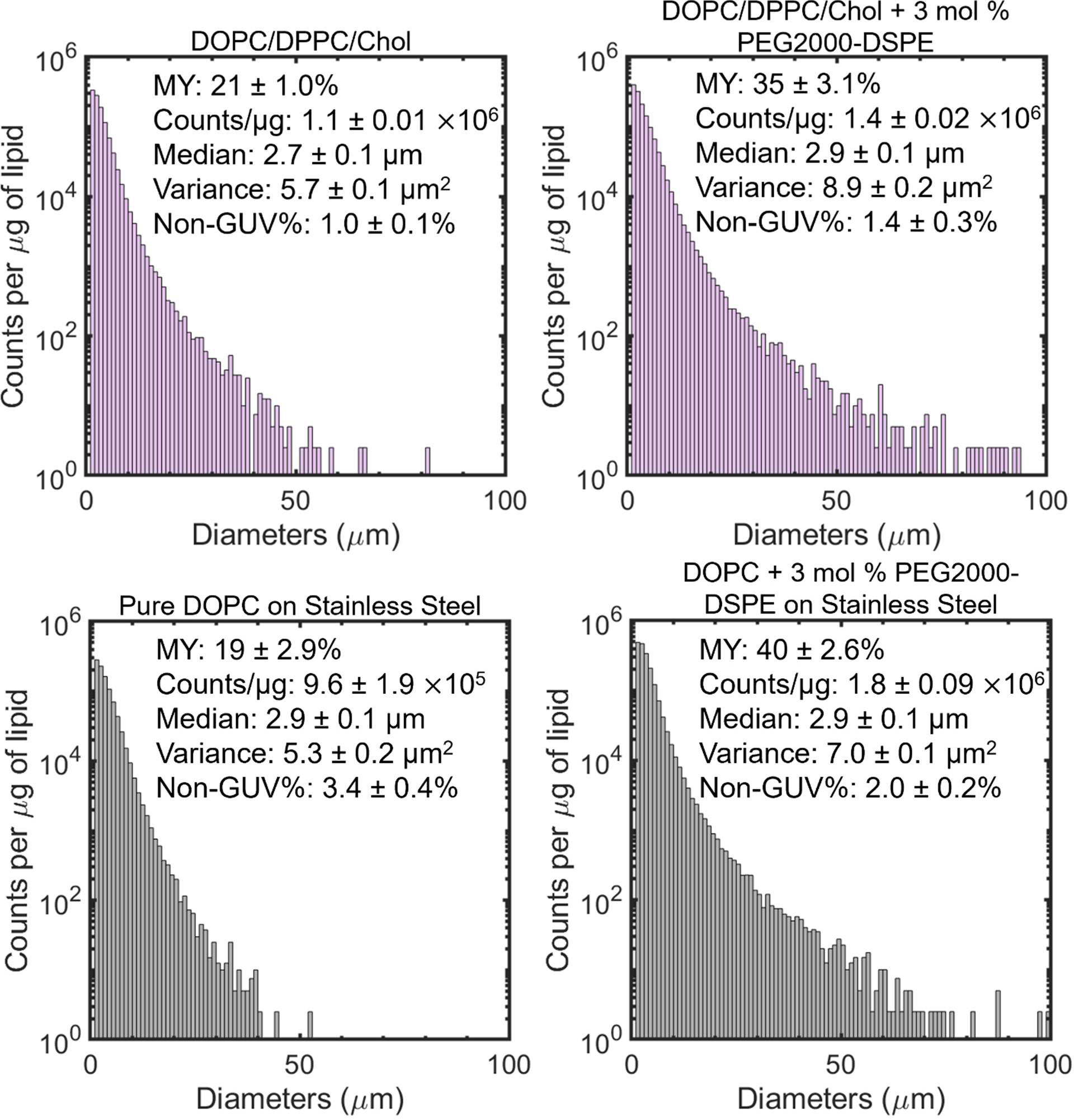
Histograms of GUV diameters of samples shown in Figure 7. Each histogram is the average of N=3 independent repeats per sample. Note the logarithmic scale on the y-axis. Bin widths are 1 µm. The molar yield (MY), total counts per µg of lipid (counts/µg), median diameter, non-GUV% and variance are shown in the text insert on the plots.

**Table S1.** Table of p-values from Student’s t-tests of the molar yields of GUVs obtained from PAPRYUS and gentle hydration on glass using DOPC with and without 3 mol % PEG2000-DSPE. *: p < 0.05, **: p < 0.01, ***: p < 0.001, NS: not significant.

| Group 1 | Group 2 | *p*-value | Significance | Interpretation |
| --- | --- | --- | --- | --- |
| PAPYRUS,<br>Pure DOPC | PAPYRUS,<br>DOPC + 3 mol<br>% PEG2000-DSPE | 0.451 | NS | 3 mol %<br>PEG2000-DSPE<br>did not result in a<br>significant change<br>in molar yield<br>using PAPYRUS |
| Gentle<br>Hydration on<br>Glass, Pure<br>DOPC | Gentle<br>Hydration on<br>Glass, DOPC +<br>3 mol %<br>PEG2000-DSPE | $5.22 \times 10^{-5}$ | *** | 3 mol %<br>PEG2000-DSPE<br>results in a<br>significant increase<br>in molar yield<br>using gentle<br>hydration on glass |

**Table S2.** The *p*-value from a Student’s t-test of the molar yields of GUVs obtained from gentle hydration on glass using DOPC with 3 mol % PEG2000-DSPE harvested at 60 minutes and 10 minutes. *: p < 0.05, **: p < 0.01, ***: p < 0.001, NS: not significant.

| Group 1 | Group 2 | *p*-value | Significance | Interpretation |
| --- | --- | --- | --- | --- |
| Gentle Hydration on Glass, 60 minute incubation | Gentle Hydration on Glass, 10 minute incubation | 0.118 | NS | Incubation for 10 minutes is not significantly different than incubation for 60 minutes |

**Table S3.** ANOVA table and table of p-values from post hoc Tukey’s HSD tests of the molar yields of GUVs obtained from gentle hydration on glass with DOPC with 0-10 mol % PEG2000-DSPE. *: p < 0.05, **: p < 0.01, ***: p < 0.001, NS: not significant.

| Source | SS | 'df' | 'MS' | 'F' | 'Prob>F' |
| --- | --- | --- | --- | --- | --- |
| Columns | 2860.373 | 5 | 572.074 | 89.746 | 4.43×-09 |
| Error | 76.492 | 12 | 6.3743 |  |  |
| Total | 2936.866 | 17 |  |  |  |

| Group 1 | Group 2 | *p*-value | Significance | Interpretation |
| --- | --- | --- | --- | --- |
| 0 mol % | 0.1 mol % | 0.119 | NS | The addition of 0.1 mol % of PEG2000-DSPE did not result in a significant change in molar yield. |
| 0 mol % | 1 mol % | 7.76×-04 | *** | The addition of 1 mol % of PEG2000-DSPE resulted in a significant change in molar yield. |
| 0 mol % | 3 mol % | 1.11×-08 | *** | The addition of 3 mol % of PEG2000-DSPE resulted in a significant change in molar yield. |
| 0 mol % | 5 mol % | 0.170 | NS | The addition of 5 mol % of PEG2000-DSPE did not result in a significant change in molar yield. |
| 0 mol % | 10 mol % | 0.439 | NS | The addition of 10 mol % of PEG2000-DSPE did not result in a significant change in molar yield. |
| 0.1 mol % | 1 mol % | 0.079 | NS | There is no significant difference between the addition of 0.1 and 1 mol % PEG2000-DSPE. |
| 0.1 mol % | 3 mol % | 9.32×-08 | *** | There is a significant difference between 0.1 and 3 mol % PEG2000-DSPE. |
| 0.1 mol % | 5 mol % | 0.999 | NS | There is no significant difference between the addition of 0.1 and 5 mol % PEG2000-DSPE. |
| 0.1 mol % | 10 mol % | 0.005 | ** | There is a significant difference between the addition of 0.1 and 10 mol % PEG2000-DSPE. |
| 1 mol % | 3 mol % | 1.47×-06 | *** | There is a significant difference between the addition of 1 and 3 mol % PEG2000-DSPE. |
| 1 mol % | 5 mol % | 0.054 | NS | There is no significant difference between the addition of 3 and 5 mol % PEG2000-DSPE. |
| 1 mol % | 10 mol % | 5.38×-05 | *** | There is a significant difference between the addition of 1 and 10 mol % PEG2000-DSPE. |
| 3 mol % | 5 mol % | 7.75×-08 | *** | There is a significant difference between the addition of 1 and 10 mol % PEG2000-DSPE. |
| 3 mol % | 10 mol % | 3.01×-09 | *** | There is a significant difference between the addition of 3 and 10 mol % PEG2000-DSPE. |
| 5 mol % | 10 mol % | 0.007 | ** | There is a significant difference between the addition of 5 and 10 mol % PEG2000-DSPE. |

**Table S4.** ANOVA table and table of p-values from post hoc Tukey’s HSD tests of the molar yields of GUVs obtained from gentle hydration on glass with pure DOPC and DOPC + 3 mol % PEG2000-DSG, DSPG, or DSPE. *: p < 0.05, **: p < 0.01, ***: p < 0.001, NS: not significant.

| Source | SS | 'df' | 'MS' | 'F' | 'Prob>F' |
| --- | --- | --- | --- | --- | --- |
| Columns | 172.811 | 3 | 57.604 | 9.741 | 0.00478 |
| Error | 47.307 | 8 | 5.913 |  |  |
| Total | 220.118 | 11 |  |  |  |

| Group 1 | Group 2 | *p*-value | Significance | Interpretation |
| --- | --- | --- | --- | --- |
| DOPC | DSPE | 0.184 | NS | Addition of 3 mol % DSPE does not result in a significant change in yield from pure DOPC |
| DOPC | PEG2000-DSG | 0.541 | NS | Addition of 3 mol % PEG2000-DSG (steric repulsion) does not result in a significant change in yield from pure DOPC |
| DOPC | DSPG | 0.091 | NS | Addition of 3 mol % DSPG (electrostatic repulsion) does not result in a significant change in yield from pure DOPC |
| DSPE | PEG2000-DSG | 0.811 | NS | Addition of DSPE and PEG2000-DSG are not significantly different from each other |
| DSPE | DSPG | $4.34 \times 10^{-3}$ | ** | Addition of DSPE and DSPG are not significantly different from each other |
| PEG2000-DSG | DSPG | $1.34 \times 10^{-2}$ | * | Addition of PEG2000-DSG and DSPG are not significantly different from each other |

**Table S5.** The *p*-value from a Student’s t-test of the molar yields of GUVs obtained from gentle hydration on glass using DOPC/DPPC/Cholesterol with and without 3 mol % PEG2000-DSPE. *: p < 0.05, **: p < 0.01, ***: p < 0.001, NS: not significant.

| Group | Group 2 | *p*-value | Significance | Interpretation |
| --- | --- | --- | --- | --- |
| DOPC/DPPC/Chol | DOPC/DPPC/Chol<br>+ 3 mol %<br>PEG2000-DSPE | $4.06 \times 10^{-3}$ | ** | Addition of 3<br>mol %<br>PEG2000-DSPE results<br>in a significant<br>increase in<br>yield |

**Table S6.** The *p*-value from a Student’s t-test of the molar yields of GUVs obtained from gentle hydration on glass using DOPC with and without 3 mol % PEG2000-DSPE on stainless steel. *: p < 0.05, **: p < 0.01, ***: p < 0.001, NS: not significant.

| Group 1 | Group 2 | *p*-value | Significance | Interpretation |
| --- | --- | --- | --- | --- |
| Pure DOPC on stainless steel | Pure DOPC + 3 mol % PEG2000-DSPE on stainless steel | $1.40 \times 10^{-3}$ | ** | Addition of 3 mol % PEG2000-DSPE results in a significant increase in yield |

## References

(1) Nair, K. S.; Bajaj, H. Advances in giant unilamellar vesicle preparation techniques and applications. Adv Colloid Interface Sci 2023, 318, 102935.

(2) York-Duran, M. J.; Godoy-Gallardo, M.; Labay, C.; Urquhart, A. J.; Andresen, T. L.; Hosta-Rigau, L. Recent advances in compartmentalized synthetic architectures as drug carriers, cell mimics and artificial organelles. Colloids Surf B Biointerfaces 2017, 152, 199–213.

(3) Has, C.; Sunthar, P. A comprehensive review on recent preparation techniques of liposomes. J Liposome Res 2020, 30, 336–365.

(4) Perrier, D. L.; Rems, L.; Boukany, P. E. Lipid vesicles in pulsed electric fields: Fundamental principles of the membrane response and its biomedical applications. Adv Colloid Interface Sci 2017, 249, 248–271.

(5) Pick, H.; Alves, A. C.; Vogel, H. Single-Vesicle Assays Using Liposomes and Cell-Derived Vesicles: From Modeling Complex Membrane Processes to Synthetic Biology and Biomedical Applications. Chem Rev 2018, 118, 8598–8654.

(6) Krinsky, N.; Kaduri, M.; Zinger, A.; Shainsky-Roitman, J.; Goldfeder, M.; Benhar, I.; Hershkovitz, D.; Schroeder, A. Synthetic Cells Synthesize Therapeutic Proteins inside Tumors. Adv Healthc Mater 2018, 7.

(7) Dhand, C.; Prabhakaran, M. P.; Beuerman, R. W.; Lakshminarayanan, R.; Dwivedi, N.; Ramakrishna, S. Role of size of drug delivery carriers for pulmonary and intravenous administration with emphasis on cancer therapeutics and lung-targeted drug delivery. RSC Adv 2014, 4, 32673–32689.

(8) Hernandez Bücher, J. E.; Staufer, O.; Ostertag, L.; Mersdorf, U.; Platzman, I.; Spatz, J. P. Bottom-up assembly of target-specific cytotoxic synthetic cells. Biomaterials 2022, 285.

(9) Parolini, L.; Mognetti, B. M.; Kotar, J.; Eiser, E.; Cicuta, P.; Di Michele, L. Volume and porosity thermal regulation in lipid mesophases by coupling mobile ligands to soft membranes. Nat Commun 2015, 6.

(10) Göpfrich, K.; Platzman, I.; Spatz, J. P. Mastering Complexity: Towards Bottom-up Construction of Multifunctional Eukaryotic Synthetic Cells. Trends Biotechnol 2018, 36, 938–951.

(11) Küchler, A.; Yoshimoto, M.; Luginbühl, S.; Mavelli, F.; Walde, P. Enzymatic reactions in confined environments. Nat Nanotechnol 2016, 11, 409–420.

(12) Cooper, A.; Girish, V.; Subramaniam, A. B. Osmotic Pressure Enables High-Yield Assembly of Giant Vesicles in Solutions of Physiological Ionic Strengths. Langmuir 2023, 39, 5579–5590.

(13) Pazzi, J.; Subramaniam, A. B. Nanoscale Curvature Promotes High Yield Spontaneous Formation of Cell-Mimetic Giant Vesicles on Nanocellulose Paper. ACS Appl Mater Interfaces 2020, 12, 56549–56561.

(14) Bangham, A. D.; Standish, M. M.; Watkins, J. C. Diffusion of univalent ions across the lamellae of swollen phospholipids. J Mol Biol 1965, 13, 238–252.

(15) Reeves, J. P.; Dowben, R. M. Formation and properties of thin-walled phospholipid vesicles. J Cell Physiol 1969, 73, 49–60.

(16) Pazzi, J.; Subramaniam, A. B. Dynamics of giant vesicle assembly from thin lipid films. J Colloid Interface Sci 2024, 661, 1033–1045.

(17) Maruyama, K.; Yuda, T.; Okamoto, A.; Kojima, S.; Suginaka, A.; Iwatsuru, M. Prolonged circulation time in vivo of large unilamellar liposomes composed of distearoyl phosphatidylcholine and cholesterol containing amphipathic poly(ethylene glycol). Biochimica et Biophysica Acta (BBA)/Lipids and Lipid Metabolism 1992, 1128, 44–49.

(18) Mori, A.; Klibanov, A. L.; Torchilin, V. P.; Huang, L. Influence of the steric barrier activity of amphipathic poly(ethyleneglycol) and ganglioside GM1 on the circulation time of liposomes and on the target binding of immunoliposomes in vivo. FEBS Lett 1991, 284, 263–266.

(19) Kumar, V.; Qin, J.; Jiang, Y.; Duncan, R. G.; Brigham, B.; Fishman, S.; Nair, J. K.; Akinc, A.; Barros, S. A.; Kasperkovitz, P. V. Shielding of Lipid Nanoparticles for siRNA Delivery: Impact on Physicochemical Properties, Cytokine Induction, and Efficacy. Mol Ther Nucleic Acids 2014, 3, e210.

(20) Higuchi, A.; Sung, T.-C.; Wang, T.; Ling, Q.-D.; Kumar, S. S.; Hsu, S.-T.; Umezawa, A. Material Design for Next-Generation mRNA Vaccines Using Lipid Nanoparticles. Polymer Reviews 2022, 63, 394–436.

(21) Cruje, C.; Chithrani, D. B. Polyethylene Glycol Functionalized Nanoparticles for Improved Cancer Treatment Article in Reviews in Nanoscience and Nanotechnology. Reviews in Nanoscience and Nanotechnology 2014, 22, 20–30.

(22) Ickenstein, L. M.; Garidel, P. Lipid-based nanoparticle formulations for small molecules and RNA drugs. Expert Opin Drug Deliv 2019, 16, 1205–1226.

(23) Large, D. E.; Abdelmessih, R. G.; Fink, E. A.; Auguste, D. T. Liposome composition in drug delivery design, synthesis, characterization, and clinical application. Adv Drug Deliv Rev 2021, 176, 113851.

(24) González-Cela-Casamayor, M. A.; López-Cano, J. J.; Bravo-Osuna, I.; Andrés-Guerrero, V.; Vicario-de-la-Torre, M.; Guzmán-Navarro, M.; Benítez-del-Castillo, J. M.; Herrero-Vanrell, R.; Molina-Martínez, I. T. Novel Osmoprotective DOPC-DMPC Liposomes Loaded with Antihypertensive Drugs as Potential Strategy for Glaucoma Treatment. Pharmaceutics 2022, 14, 1405.

(25) Tam, A.; Kulkarni, J.; An, K.; Li, L.; Dorscheid, D.; Singhera, G.; Bernatchez, P.; Reid, G.; Chan, K.; Witzigmann, D.;, et al. Lipid nanoparticle formulations for optimal RNA-based topical delivery to murine airways. European Journal of Pharmaceutical Sciences 2022, 176, 106234.

(26) Wagner, M. J.; Mitra, R.; Mcarthur, M. J.; Baze, W.; Barnhart, K.; Wu, S. Y.; Rodriguez-Aguayo, C.; Zhang, X.; Coleman, R. L.; Lopez-Berestein, G.;, et al. Preclinical mammalian safety studies of EPHARNA (DOPC nanoliposomal EphA2-targeted siRNA). Mol Cancer Ther 2017, 16, 1114.

(27) Shi, D.; Beasock, D.; Fessler, A.; Szebeni, J.; Ljubimova, J. Y.; Afonin, K. A.; Dobrovolskaia, M. A. To PEGylate or not to PEGylate: Immunological properties of nanomedicine’s most popular component, polyethylene glycol and its alternatives. Adv Drug Deliv Rev 2022, 180, 114079.

(28) Dilliard, S. A.; Siegwart, D. J. Passive, active and endogenous organ-targeted lipid and polymer nanoparticles for delivery of genetic drugs. Nature Reviews Materials 2023 8:4 2023, 8, 282–300.

(29) Hald Albertsen, C.; Kulkarni J. A.; Witzigmann D.; Lind M.; Petersson K.; Simonsen, J. B. The role of lipid components in lipid nanoparticles for vaccines and gene therapy. Adv Drug Deliv Rev 2022, 188, 114416.

(30) Suk, J. S.; Xu, Q.; Kim, N.; Hanes, J.; Ensign, L. M. PEGylation as a strategy for improving nanoparticle-based drug and gene delivery. Adv Drug Deliv Rev 2016, 99, 28– 51.

(31) Needham, D.; Hristova, K.; Mcintosh, T. J.; Dewhirst, M.; Wu, N.; Lasic, D. D. Polymer-grafted liposomes: Physical basis for the “stealth” property. J Liposome Res 1992, 2, 411– 430.

(32) Efremova, N. V.; Bondurant, B.; O’Brien, D. F.; Leckband, D. E. Measurements of interbilayer forces and protein adsorption on uncharged lipid bilayers displaying poly(ethylene glycol) chains. Biochemistry 2000, 39, 3441–3451.

(33) Kenworthy, A. K.; Simon, S. A.; Mcintosh, T. J. Structure and Phase Behavior of Lipid Suspensions Containing Phospholipids with Covalently Attached Poly(ethylene glycol). Biophys J 1995, 68.

(34) Kenworthy, A. K.; Hristova, K.; Needham, D.; McIntosh, T. J. Range and magnitude of the steric pressure between bilayers containing phospholipids with covalently attached poly(ethylene glycol). Biophys J 1995, 68, 1921–1936.

(35) Kuhl, T. L.; Leckband, D. E.; Lasic, D. D.; Israelachvili, J. N. Modulation of Interaction Forces Between Bilayers Exposing Short-Chained Ethylene Oxide Headgroups. Biophys J 1994, 66, 1479–1488.

(36) Needham, D.; Mclatosh, T. J.; Lasic, D. D. Repulsive interactions and mechanical stability of polymer-grafted lipid membranes. Biochimiea el Biophysica Acta 1992, 1108, 40–48.

(37) Hristova, K.; Needham, D. Phase Behavior of a Lipid/Polymer-Lipid Mixture in Aqueous Medium. Macromolecules 1995, 28, 991–1002.

(38) Belsito, S.; Bartucci, R.; Montesano, G.; Marsh, D.; Sportelli, L. Molecular and mesoscopic properties of hydrophilic polymer-grafted phospholipids mixed with phosphatidylcholine in aqueous dispersion: Interaction of dipalmitoyl N-poly(ethylene glycol)phosphatidylethanolamine with dipalmitoylphosphatidylcholine studied by spectrophotometry and spin-label electron spin resonance. Biophys J 2000, 78, 1420– 1430.

(39) Marsh, D.; Bartucci, R.; Sportelli, L. Lipid membranes with grafted polymers: Physicochemical aspects. Biochim Biophys Acta Biomembr 2003, 1615, 33–59.

(40) Viitala, L.; Pajari, S.; Gentile, L.; Määttä, J.; Gubitosi, M.; Deska, J.; Sammalkorpi, M.; Olsson, U.; Murtomäki, L. Shape and Phase Transitions in a PEGylated Phospholipid System. Langmuir 2019, 35, 3999–4010.

(41) Hristova, K.; Needham, D. The Influence of Polymer-Grafted Lipids on the Physical Properties of Lipid Bilayers: A Theoretical Study. J Colloid Interface Sci 1994, 168, 302–314.

(42) Marsh, D. Elastic constants of polymer-grafted lipid membranes. Biophys J 2001, 81, 2154–2162.

(43) Lasic, D. D.; Joannic, R.; Keller, B. C.; Frederik, P. M.; Auvray, L. Spontaneous vesiculation. Adv Colloid Interface Sci 2001, 337–349.

(44) Mahendra, A.; James, H. P.; Jadhav, S. PEG-grafted phospholipids in vesicles: Effect of PEG chain length and concentration on mechanical properties. Chem Phys Lipids 2019, 218, 47–56.

(45) Bivas, I.; Vitkova, V.; Mitov, M. D.; Winterhalter, M.; Alargova, R. G.; Méléard, P.; Bothorel, P. Mechanical Properties of Lipid Bilayers Containing Grafted Lipids. In Giant Vesicles: Perspectives in Supramolecular Chemistry; 2000.

(46) Silvander, M.; Johnsson, M.; Edwards, K. Effects of PEG-lipids on permeability of phosphatidylcholine/cholesterol liposomes in buffer and in human serum. Chem Phys Lipids 1998, 97, 15–26.

(47) Hashizaki, K.; Taguchi, H.; Itoh, C.; Sakai, H.; Abe, M.; Saito, Y.; Ogawa, N. Effects of Poly(ethylene glycol) (PEG) Chain Length of PEG-Lipid on the Permeability of Liposomal Bilayer Membranes. Chem. Pharm. Bull. 2003, 51, 815–820.

(48) Lee, H.; Larson, R. G. Adsorption of Plasma Proteins onto PEGylated Lipid Bilayers: The Effect of PEG Size and Grafting Density. Biomacromolecules 2016, 17, 1757–1765.

(49) Giakoumatos, E. C.; Gascoigne, L.; Gumí-Audenis, B.; García, Á. G.; Tuinier, R.; Voets, I. K. Impact of poly(ethylene glycol) functionalized lipids on ordering and fluidity of colloid supported lipid bilayers. Soft Matter 2022, 18, 7569–7578.

(50) Stepniewski, M.; Pasenkiewicz-Gierula, M.; Og, T. R.; Danne, R.; Orlowski, A.; Karttunen, M.; Urtti, A.; Yliperttula, M.; Vuorimaa, E.; Bunker, A. Study of PEGylated Lipid Layers as a Model for PEGylated Liposome Surfaces: Molecular Dynamics Simulation and Langmuir Monolayer Studies. Langmuir 2011, 27, 7788–7798.

(51) Bartucci, R.; Pantusa, M.; Marsh, D.; Sportelli, L. Interaction of human serum albumin with membranes containing polymer-grafted lipids: spin-label ESR studies in the mushroom and brush regimes. Biochimica et Biophysica Acta (BBA) - Biomembranes 2002, 1564, 237–242.

(52) Shohda, K.; Takahashi, K.; Suyama, A. A method of gentle hydration to prepare oil-free giant unilamellar vesicles that can confine enzymatic reactions. Biochem Biophys Rep 2015, 3, 76–82.

(53) Yamashita, Y.; Oka, M.; Tanaka, T.; Yamazaki, M. A new method for the preparation of giant liposomes in high salt concentrations and growth of protein microcrystals in them. Biochim Biophys Acta Biomembr 2002, 1561, 129–134.

(54) Tsai, F. C.; Stuhrmann, B.; Koenderink, G. H. Encapsulation of active cytoskeletal protein networks in cell-sized liposomes. Langmuir 2011, 27, 10061–10071.

(55) Schindelin, J.; Arganda-Carreras, I.; Frise, E.; Kaynig, V.; Longair, M.; Pietzsch, T.; Preibisch, S.; Rueden, C.; Saalfeld, S.; Schmid, B.; et al. Fiji: An open-source platform for biological-image analysis. Nat Methods 2012, 9, 676–682.

)56) Israelachvili, J. N. *Intermolecular and Surface Forces*, 3rd ed.; Academic Press: Waltham, MA, 2011; Vol. 3.

(57) Bekmurzayeva, A.; Duncanson, W. J.; Azevedo, H. S.; Kanayeva, D. Surface modification of stainless steel for biomedical applications: Revisiting a century-old material. Materials Science and Engineering: C 2018, 93, 1073–1089.

## Supporting reference

1. Cooper, A., Girish, V. & Subramaniam, A.B. Osmotic Pressure Enables High-Yield Assembly of Giant Vesicles in Solutions of Physiological Ionic Strengths. Langmuir 39, 5579–5590 (2023).

